# Satellite glial GPR37L1 regulates maresin and potassium channel signaling for pain control

**DOI:** 10.1101/2023.12.03.569787

**Authors:** Sangsu Bang, Changyu Jiang, Jing Xu, Sharat Chandra, Aidan McGinnis, Xin Luo, Qianru He, Yize Li, Zilong Wang, Xiang Ao, Marc Parisien, Lorenna Oliveira Fernandes de Araujo, Sahel Jahangiri Esfahan, Qin Zhang, Raquel Tonello, Temugin Berta, Luda Diatchenko, Ru-Rong Ji

## Abstract

G protein coupled receptor 37-like 1 (GPR37L1) is an orphan GPCR and its function remains largely unknown. Here we report that GPR37L1 transcript is highly expressed compared to all known GPCRs in mouse and human dorsal root ganglia (DRGs) and selectively expressed in satellite glial cells (SGCs). Peripheral neuropathy following diabetes and chemotherapy by streptozotocin and paclitaxel resulted in downregulations of surface GPR37L1 in mouse and human DRGs. Transgenic mice with *Gpr37l1* deficiency exhibited impaired resolution of neuropathic pain symptom (mechanical allodynia), whereas overexpression of *Gpr37l1* in mouse DRGs can reverse neuropathic pain. Notably, GPR37L1 is co-expressed and coupled with potassium channels in SGCs. We found striking species differences in potassium channel expression in SGCs, with predominant expression of KCNJ10 and KCNJ3 in mouse and human SGCs, respectively. GPR37L1 regulates the surface expression and function of KCNJ10 and KCNJ3. We identified the pro-resolving lipid mediator maresin 1 (MaR1) as a GPR37L1 ligand. MaR1 increases KCNJ10/KCNJ3-mediated potassium influx in SGCs via GPR37L1. MaR1 protected chemotherapy-induced suppression of KCNJ13/KCNJ10 expression and function in SGCs. Finally, genetic analysis revealed that the *GPR37L1-E296K* variant is associated with increased chronic pain risk by destabilizing the protein. Thus, GPR37L1 in SGCs offers a new target for neuropathy protection and pain control.

## Introduction

Satellite glial cells (SGCs) reside in the peripheral nervous system (PNS), where they wrap around neuronal cell bodies and form a complete envelope, allowing for close neuron-SGC interactions in the dorsal root ganglia (DRGs) and trigeminal ganglia (1, 2). SGCs express high levels of the inwardly rectifying K^+^ channel KCNJ10 (also known as Kir4.1) in mouse sensory ganglia, enabling them to control perineural potassium homeostasis and neuronal excitability (2, 3). Increasing evidence indicates that SGCs participate in the generation and maintenance of chronic pain (4-6). Especially, KCNJ10 is downregulated under pathological pain conditions (7), and the knockdown of KCNJ10 expression in SGCs is sufficient to induce pain hypersensitivity (3). However, it is unclear how KCNJ10 expression is regulated in SGCs and disease conditions. It is generally believed that SGCs promote pain by ATP signaling or releasing pro-inflammatory cytokines, such as TNF-α and IL-1β, which can drive hyperexcitability of surrounding sensory neurons (2, 8, 9).

G-protein coupled receptor 37-like 1 (GPR37L1) is an orphan GPCR. Prosaposin and prosaposin-derived peptides such as TX-14 have been proposed as ligands of GPR37 and GPR37L1, demonstrating neuroprotective and glioprotective effects. However, the precise binding sites of these ligands on GPR37L1 remain elusive (10, 11). Studies have shown that GPR37L1 exhibits constitutive activities regulated by protease cleavage (12) and remains unliganded (13). *GPR37L1* mutations have been implicated in neurological diseases, such as seizure susceptibility (14-16). Single-cell analysis revealed that the *Gpr37l1* transcript is highly enriched in astrocytes and SGCs (17-22). Despite this, the exact role of GPR37L1 in SGCs is unclear.

Specialized pro-resolving mediators (SPMs), such as resolvins, protectins, and maresins, are biosynthesized from omega-3 unsaturated fatty acids (e.g., DHA), and synthetic SPMs exhibit potent pro-resolution, anti-inflammation, and analgesic actions in various animal models via GPCR activation (23-27). We previously identified neuroprotectin D1 (NPD1) as a ligand for GPR37 (28). In this study, we identified DHA-derived maresin 1 (MaR1) as a potential ligand of GPR37L1. We found that MaR1 interacts with and binds GPR37L1. MaR1 potently attenuated neuropathic pain and further increased KCNJ10-mediated K^+^ currents in SGCs, in a GPR37L1-dependent manner. We also revealed a distinct expression of KCNJ3 (Kir3.1/GIRK1) in humans but not in mouse SGCs and demonstrated that MaR1 increased the KCNJ3 activity in human SGCs. Finally, we identified novel *GPR37L1* variants that are associated with chronic pain in humans.

## Results

### *Gpr37l1* and *GPR37L1* are highly expressed in SGCs of mouse and human DRG

We conducted RNA sequencing from DRGs of non-injured naive mice and detected a total of 321 GPCRs and 92 orphan GPCRs with a cutoff value of >1 (Supplemental Table 1A). Figure 1A and B are lists of the top ten expressed GPCR transcripts in both categories. We found that *Gpr37l1* ranks among the ten most highly expressed GPCR mRNAs (Figure 1A; *n* = 3). Furthermore, *Gpr37l1* stands out as the most highly expressed orphan GPCR mRNA in mouse DRGs (Figure 1B; Supplemental Table 1B*, n* = 3). Our investigation then extended to examine GPCR mRNA expression levels in human DRGs, utilizing a microarray database encompassing a total of 332 GPCR and 73 orphan GPCR transcripts (Supplemental Table 2A; *n* = 214). Figure 1C and 1D show the ten most highly expressed GPCR transcripts in both categories. We found that *GPR37L1* ranks among the top ten GPCR mRNAs (Figure 1C, Supplemental Table 2A) and the top five orphan GPCR mRNAs in human DRGs (Figure 1D, Supplemental Table 2B). Therefore, *Gpr37l1* and *GPR37L1* mRNAs are highly expressed by both mouse and human DRGs.

**Figure 1.**
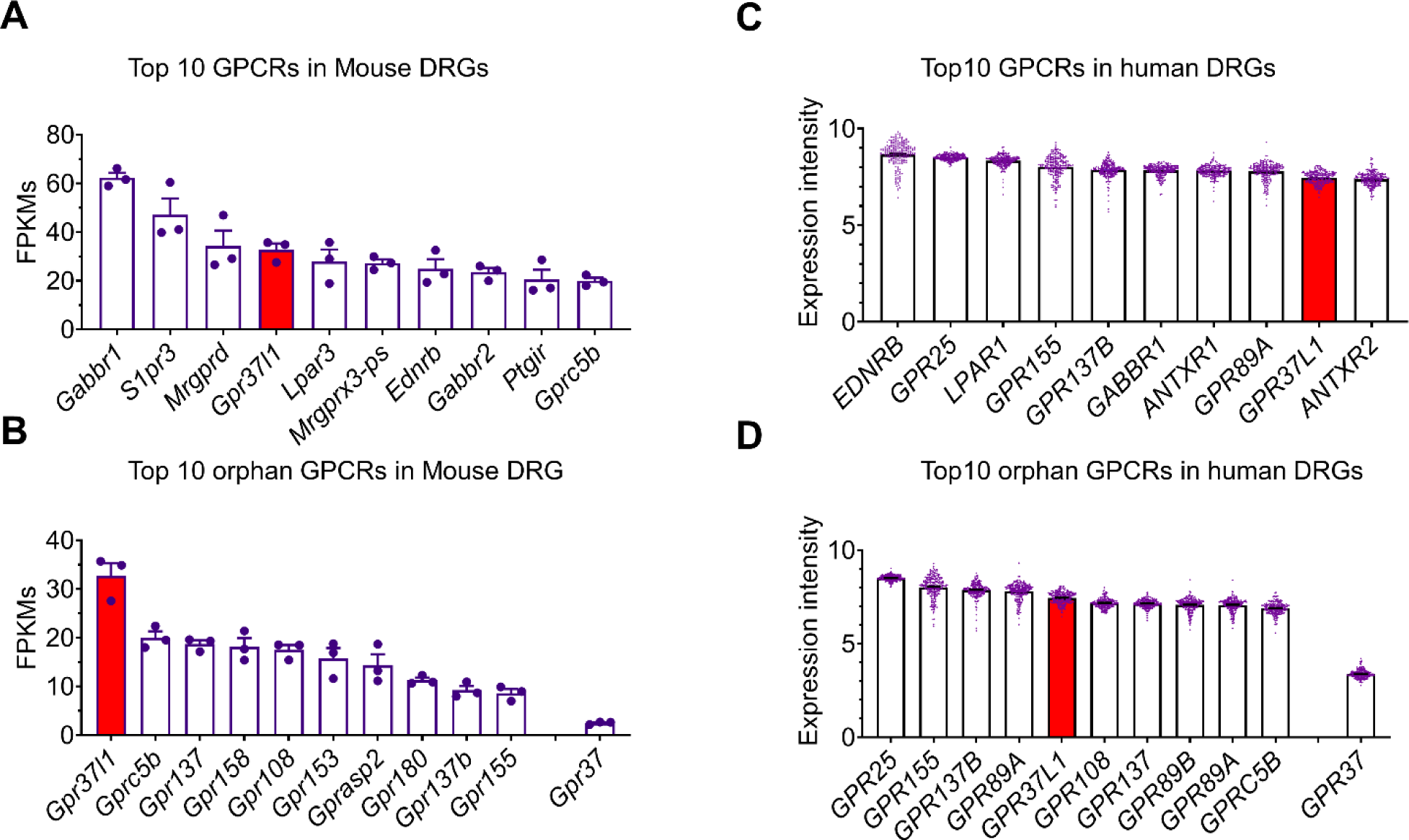
*Gpr37l1* and *GPR37L1* transcripts are highly expressed in mouse and human DRGs. (**A, B**) Highly expressed GPCR transcripts in mouse DRGs. RNAseq shows mRNA expression levels (FPKM) of the top 10 GPCR transcripts (A, *n* = 3) and the top 10 orphan GPCR transcripts (B, *n* = 3). *Gpr37l1* expression is highlighted in red bars. Note that the *Gpr37* expression is much lower than *Gpr37l1*. (**C, D**) Highly expressed GPCR transcripts in human DRGs. Normalized microarray shows mRNA expression levels (intensity) of the top 10 GPCR transcripts (C, *n* = 214) and the top 10 orphan GPCR transcripts (D, *n* = 214). *GRP37L1* expression is highlighted in red bars. *GPR37* expression is included for comparison. Data are expressed as mean ± SEM.

Single-cell RNA sequencing (RNAseq) has revealed selective expression of *Gpr37l1* mRNA in mouse satellite glial cells (SGCs) of mouse DRG (17, 19-21, 29, 30) (Supplemental Figure 1A). Next, we investigated *Gpr37l1* mRNA and GPR37L1 protein expression in mouse DRGs using RNAscope in situ hybridization (ISH), immunohistochemistry (IHC), western blotting, and flow cytometry in both wildtype (WT) and *Gpr37l1* mutant mice (Supplemental Figure 2, A-D). ISH analysis revealed *Gpr37l1* expression in SGCs but not neurons of mouse DRG (Figure 2A). Additionally, SGCs of trigeminal ganglia and nodose ganglia expressed *Gpr37l1* (Supplemental Figure 1, B-C). *Gpr37l1* expression was absent in *Gpr37l1*^-/-^ (KO) mice, validating the specificity of the RNAscope probe (Supplemental Figure 1D). Notably, *Gpr37*, a close family member of *Gpr37l1*, is expressed by neurons but not by SGCs (Supplemental Figure 1E) in mouse DRGs. Double staining via IHC showed co-localization of GPR37L1 with FABP7, a cellular marker for SGCs (19). Notably, GPR37L1 staining forms a ring surrounding DRG neurons (Figure 2B). Western blot analysis detected several forms of GPR37L1 in mouse DRGs, a full-length GPR37L1 at ∼65 kDa (glycosylated form), and a ∼50 kDa band (non-glycosylated form), as well as truncated/cleaved forms of GPR37L1 at 37, and 25 kDa (Figure 2C). These bands were substantially reduced in the heterozygote (*Gpr37L1*^+/-^) mice and completely abolished in homozygote (*Gpr37l1*^-/-^) mice (Figure 2C). Flow cytometry analysis showed co-localization of GPR37L1 with ∼80% of GLAST^+^ cells in WT DRGs (Supplemental Figure 2A and 2B). GPR37L1 expression was abolished in *Gpr37l1*^-/-^ mice (*P*< 0.0001, *n =* 5 mice/group, Supplemental Figure 2B) and also significantly decreased in mice treated with intraganglionic (IG) injection of *Gpr37l1* siRNA (*P*< 0.001, *n =* 3, Supplemental Figure 2C).

**Figure 2.**
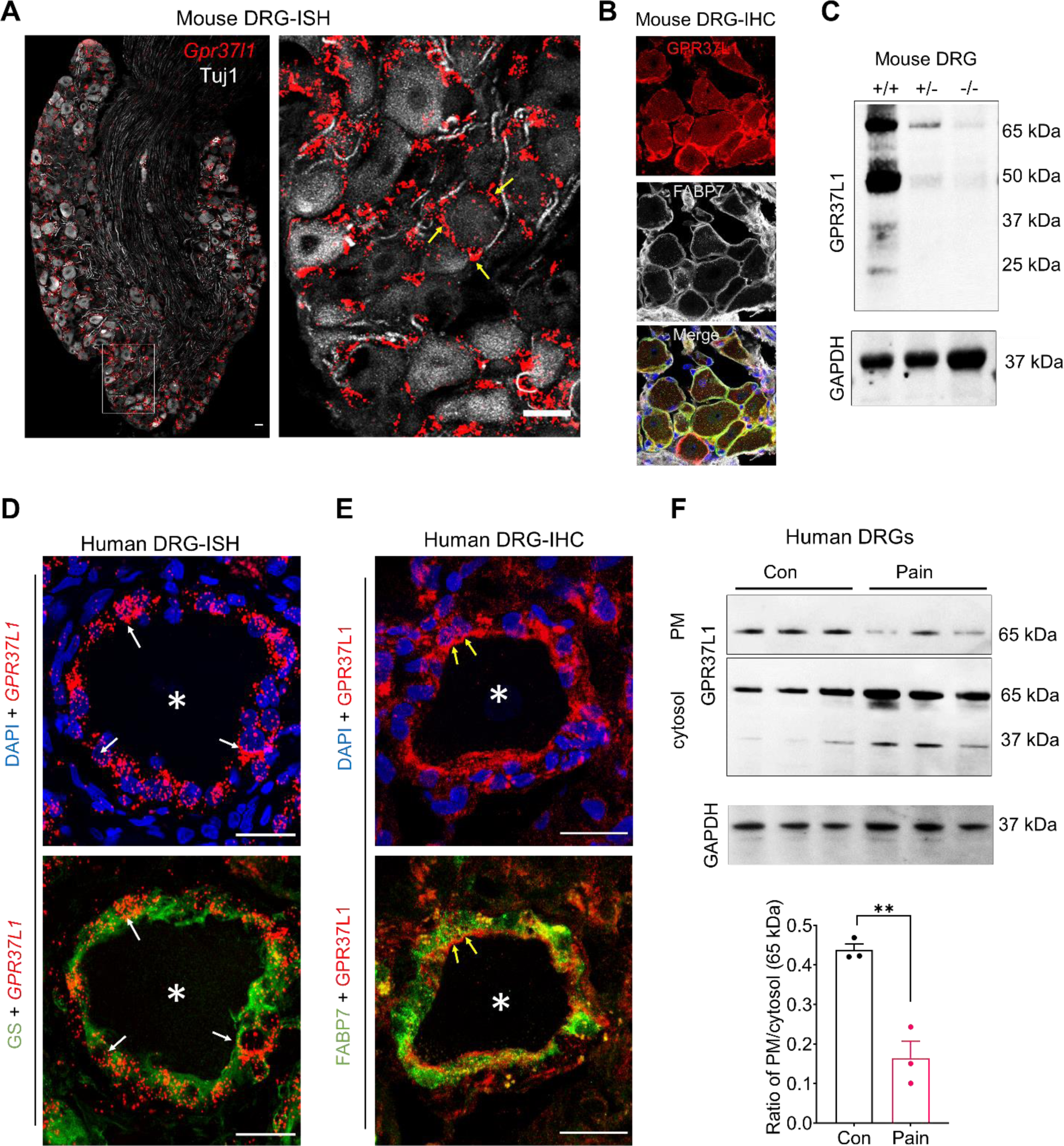
Mouse and Human SGCs express *Gpr37l1*/*GPR37L1* mRNA and GPR37L1 protein. (**A**) Double staining of RNAscope in situ hybridization (ISH for *Gpr37l1*) and immunohistochemistry (IHC for Tuj1) show non-overlapping expression of *Gpr37l1* mRNA (red) and Tuj1 (white) in mouse DRGs. Right, enlarged image from the box in the left panel. Yellow arrows indicate *Gpr37l1*^+^ cells surrounding DRG neurons. Scale = 25 µm. **(B)** Double IHC staining shows co-localization of GPR37L1 (red) with FABP7 (white). Scale bars, 25 µm. (**C**) Western blot showing GPR37L1 expression in mouse DRGs of *Gpr37l1* ^+/+^*, Gpr37l1* ^+/-^*, and Gpr37l1* ^-/-^ mice. GAPDH was included as a loading control. (**D**) Double staining of ISH (*GPR37L1,* red) and IHC (glutamine synthetase, GS, green) shows co-localization of *GPR37L1* mRNA and GS in human SGCs. **(E)** Double staining of IHC for GPR37L1 (red) and FABP7 (green) shows heavy co-localization of GPR37L1 and FABP7 in human SGCs. Note that GPR37L1 is enriched on the inner side of SGCs in close contact with neurons. * indicates a neuron and arrows indicate SGCs surrounding the neurons. Scale bars, 25 µm. **(F)** Top, western blots showing plasma membrane (PM) and cytosol fractions of GPR37L1 and GAPDH loading control in human DRGs of neuropathic pain patients and controls (Con, *n* = 3). Bottom, the ratio of PM/cytosol GPR37L1 expression in human DRGs of control and neuropathic pain patients. Data are expressed as mean ± SEM and analyzed by t-test. \*\**P*<0.05, *n* = 3, unpaired student’s t-test.

We also conducted ISH, IHC, and western blotting to examine GPR37L1 expression in human DRG tissues. Double staining of ISH and IHC revealed that the *GPR37L1* transcript is specifically expressed in human SGCs that co-express SGC marker glutamine synthetase (GS) (2) (Figure 2D). Furthermore, double IHC staining showed heavy co-localization of GPR37L1 with FABP7. Intriguingly, GPR37L1 expression on SGCs appears to be polarized: it is highly visible on the inner side of the SGC that is in close contact with neurons (Figure 2E). This unique subcellular expression provided an anatomical substrate for GPR37L1 to mediate neuron-glial interaction.

To further the translational relevance of this study, we analyzed human DRG samples from patients with diabetic peripheral neuropathy (DPN). Western blot analysis revealed a down-regulation of plasma membrane GPR37L1 (GPR37L1-PM) but an upregulation of intracellular cytosol GPR37L1 (GPR37L1-IC), leading to a significant change in PM/IC ratio (*P*<0.05, Figure 2F, *n* = 3). This result suggests that GPR37L1 is regulated in painful disease conditions such as diabetic neuropathy.

### GPR37L1 is protective against PTX and STZ-induced pain

To evaluate the contribution of GPR37L1 to neuropathic pain, we generated two animal models by systemic injection of streptozotocin (STZ), a diabetes-inducing toxin, and paclitaxel (PTX), a chemotherapy drug. We tested the time course of STZ and PTX-induced mechanical allodynia, a cardinal feature of neuropathic pain in these mouse models in three genotypes: *Gpr37l1*^+/+^, *Gpr37l1*^+/-^, and *Gpr37l1*^-/-^. Notably, the baseline pain sensitivity, including mechanical, heat, and cold sensitivity, did not differ among the three genotypes (Supplemental Figure 3A-3C). A low dose of STZ (75 mg/kg) evoked rapid mechanical allodynia in 7 days, and this mechanical pain resolved on Day 35 (Figure 3A). Interestingly, *Gpr37l1*^-/-^ mice failed to resolve on Day 35 and Day 42, compared to *Gpr37l1*^+/+^ mice (*P*<0.05 on Day 35 and Day 42, Figure 3A). A single injection of PTX (6 mg/kg) also elicited profound mechanical allodynia in 7 days, and this allodynia resolved on Day 35 (Figure 3B). Notably, *Gpr37l1*^-/-^ mice showed no sign of resolution on Day 35 and Day 42, compared to *Gpr37l1*^+/+^ mice (*P*<0.01 on Day 35, *P*<0.0001 on Day 42, Figure 3B). Taken together these findings suggest a protective role of GPR37L1 against STZ and PTX-induced pain. Consistent with this notion, the STZ-induced pain model was associated with a significant reduction of surface expression of GPR37L1 in the plasma membrane (PM) of mouse DRGs collected 28 days after the STZ injection (*P*<0.05, Figure 3C, D).

**Figure 3.**
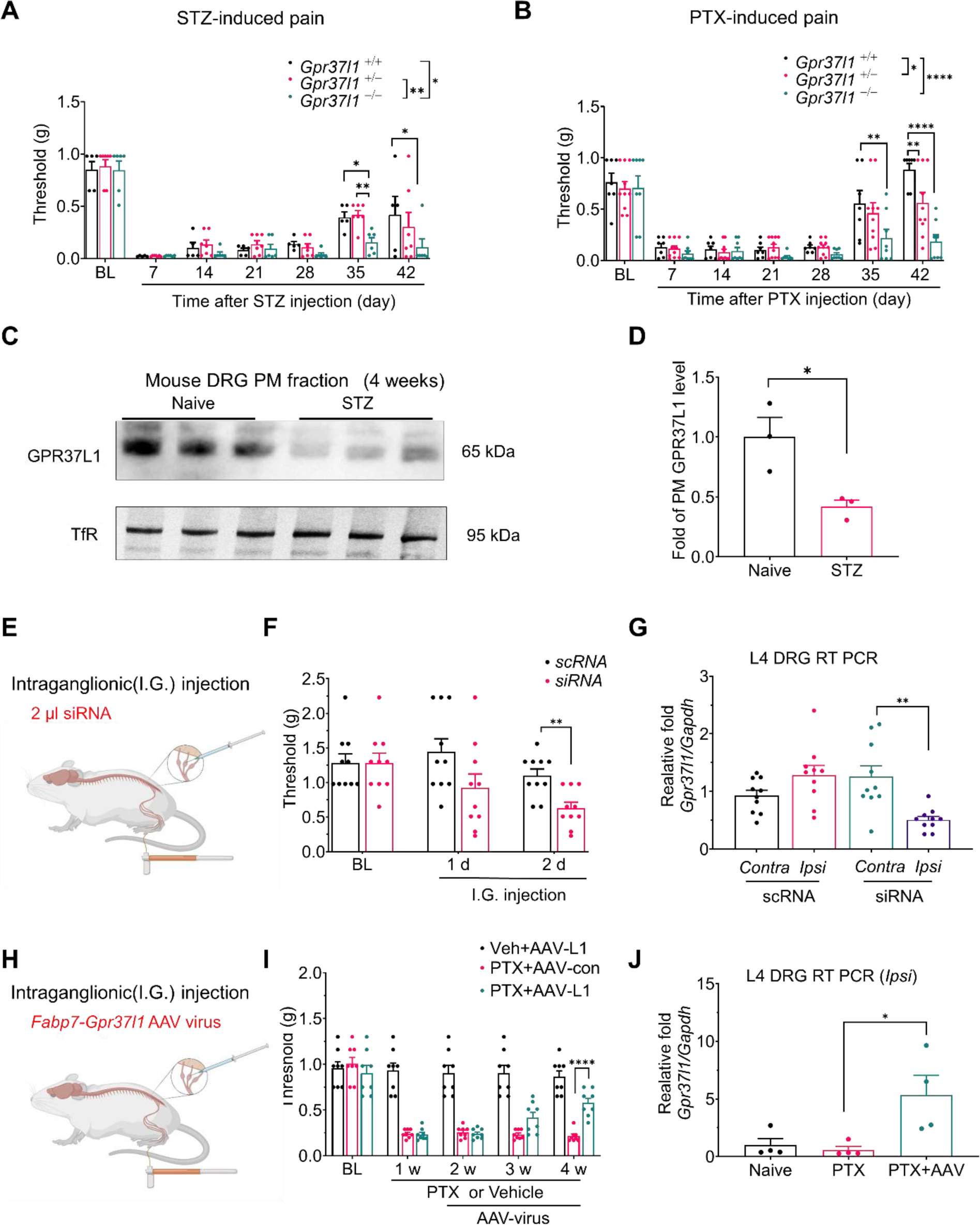
GPR37L1 is dysregulated in pain and protects against neuropathic pain. (**A, B**) Neuropathic pain (mechanical allodynia) induced by streptozotocin (75 mg/kg STZ, A) and paclitaxel (6 mg/kg, PTX, B) in wild-type and *Gpr37l1* mutant mice. (**A**) Time course of STZ-induced mechanical allodynia in *Gpr37l1* ^+/+^ mice (*n =* 5)*, Gpr37l1* ^+/-^ mice (*n =* 7)*, and Gpr37l1* ^-/-^ mice (*n =* 6). (**B**) Time course of PTX-induced mechanical allodynia in *Gpr37l1* ^+/+^ mice (*n =* 7)*, Gpr37l1* ^+/-^ mice (*n =* 10)*, and Gpr37l1* ^-/-^ mice (*n =* 8). (**C, D**) GPR37L1 expression in the plasma membrane (PM) fraction of DRG tissues of control animals (*n* = 3) and animals 4 weeks after STZ treatment (*n* = 3). Transferrin (TfR) was used as a loading control for surface proteins. (**D**) Quantification of GPR37L1 as fold change from control. (**E-G**) Unilateral intraganglionic (I.G.) microinjection of *Gpr37l1*-targeting siRNA reduces *Gpr37l1* expression and induces persistent mechanical allodynia in naïve animals. **(E**) Schematic of unilateral I.G. microinjection (2 μl) of *Gpr37l1*-targeting siRNA (siRNA) and scrambled control RNA (scRNA) in the L4 and L5 DRGs, followed by von Frey testing on day 1 and day 2 and subsequent tissue collection for quantitative RT-PCR (qPCR) analysis. **(F)** Mechanical allodynia is induced by *siRNA* (*n* = 10 mice) but not *scRNA* (*n* = 10 mice). (**G**) qPCR analyses showing expression of *Gpr37l1* in L4-L5 DRGs from the ipsilateral (*Ipsi*) and contralateral (*Contra*) sides 2 days after unilateral injection of *siRNA* (*n* = 10 mice) and *scRNA* (*n* = 10 mice). (**H-J**) Unilateral I.G. microinjection of *Fabp7*-*Gpr37l1* or Fabp7-mock AAV virus rescued *Gpr37l1* expression and reduced persistent mechanical allodynia in CIPN mice (6 mg/kg PTX). **(H**) Schematic of intraganglionic unilateral microinjection (2 μl) of *Fabp7* promoter *Gpr37l1* expression AAV9 virus (AAV-L1) and Mock control virus (AAV-Con) in the L4 and L5 DRGs, given one week after PTX, followed by von Frey testing and subsequent tissue collection for quantitative RT-PCR (qPCR) analysis. **(I)** PTX-mediated mechanical allodynia is reduced by AAV-L1 application (*n* = 8 mice) but not AAV-Con (*n* = 8 mice). Note that AAV-L1 injection in control mice without PTX treatment has no effects on mechanical pain. (**J**) qPCR analyses showing expression of *Gpr37l1* in ipsilateral L4 DRGs (*Ipsi, n* = 4 mice) 4 weeks after unilateral virus injection in naïve and PTX mice. Data are expressed as mean ± SEM and statistically analyzed by Two-Way ANOVA with Tukey’s post-hoc test (A, B, and I) or Bonferroni’s posthoc test (F), One-Way ANOVA with Tukey’s post-hoc test (G and J), and two-tailed t-test (D). \**P*<0.05, ** *P*<0.01, \*\*\*\**P*<0.0001.

To examine whether down-regulation of GPR37L1 is sufficient to produce pain, we conducted intra-ganglionic microinjection of *Gpr37l1*-targeting siRNA (siRNA) or control scramble RNA (scRNA) to the L4 and L5 DRGs (4 μg in 2 μl per injection, Figure 3E). We found that this siRNA treatment was sufficient to induce mechanical allodynia within 2 days (Figure 3F, Supplemental Figure 3D), with mild effects on thermal sensitivity (Supplemental Figure 3, E and F). Compared to scRNA, siRNA-treated mice had a ∼ 60% reduction in *Gpr37l1* mRNA levels in the L4-L5 DRGs two days after the injection (*P*<0.001, vs. scRNA, Figure 3G). A similar reduction in GPR37L1 expression was validated by flow cytometry (Supplemental Figure 2C). This finding suggests that a partial loss of GPR37L1 in L4-L5 DRGs is sufficient to drive pain.

Next, we tested whether an up-regulation of GPR37L1 in DRG SGCs is sufficient to rescue neuropathic pain. To this end, we conducted intra-ganglionic microinjection of SGC-targeting *Gpr37l1-* AAV9 virus (with *Fabp7* promoter) and control AAV9 virus (1 x 10^12^ CFU in 2 μl) to the L4 and L5 DRGs, given one week after PTX injection (Figure 3H). The *Gpr37l1*-AAV9 treatment resulted in a delayed effect on mechanical allodynia: *Gpr37l1*-AAV9 treatment reversed mechanical allodynia, caused by PTX at 4 weeks (*P*<0.0001, vs. control AAV9, Figure 3I). As expected, *Gpr37l1*-AAV9 treatment also resulted in a significant increase in *Gpr37l1* mRNA levels in the L4 DRGs at 4 weeks post-PTX injection (*P*<0.05, Figure 3J). Together, both loss-of-function and gain-of-function experiments support a protective action of GPR37L1 against neuropathic pain.

### MaR1 forms a binding interaction with GPR37L1

Specialized pro-resolving mediators (SPMs), such as resolvins, protectins, and maresins, are biosynthesized from omega-3 unsaturated fatty acids, such as docosahexaenoic acid (DHA) and demonstrate potent analgesic actions in various animal models via activation of specific GPCRs (23-27). We previously identified neuroprotectin D1 (NPD1) as a ligand for GPR37 (28). Since GPR37L1 is a close family member of GPR37, we speculated that SPMs may also activate GPR37L1. We tested different families of SPMs, including D-series resolvins (RvD1, RvD2, and RvD5), E-series resolvin (RvE1), NPD1, and maresin 1 (MaR1), as well as the SPM precursors DHA and EPA (eicosapentaenoic acid). This was accomplished by using a lipid overlay assay to detect binding partners of GPR37L1, which we established by transfecting HEK cells with *GPR37L1* containing a FLAG tag (Figure 4A). Among all the lipid mediators we tested, only MaR1 showed a specific binding signal to GPR37L1; no signal was detected following MOCK transfection (Figure 4B). To confirm whether native GPR37L1 expressed by mouse DRGs would also interact with MaR1, we performed a lipid overlay assay using DRG lysates from WT or *Gpr37l1*^-/-^ mice (Figure 4C). We coated PVDF membranes with MaR1 (0.01, 0.1, 1, and 10 nM) and observed specific binding of 10 nM MaR1 in DRG samples from WT but not from *Gpr37l1*^-/-^ mice (*P*<0.05, WT vs. KO, *n =* 3, Figure 4C). Furthermore, we performed a lipid-coated bead pull-down assay to test whether MaR1 would interact with the GPR37L1 protein. We coated beads with MaR1, RvD1, RvD2, NPD1, and DHA and found strong GPR37L1 binding to MaR1 (Supplemental Figure 4, A-C). These results suggested a direct binding of MaR1 to GPR37L1.

**Figure. 4.**
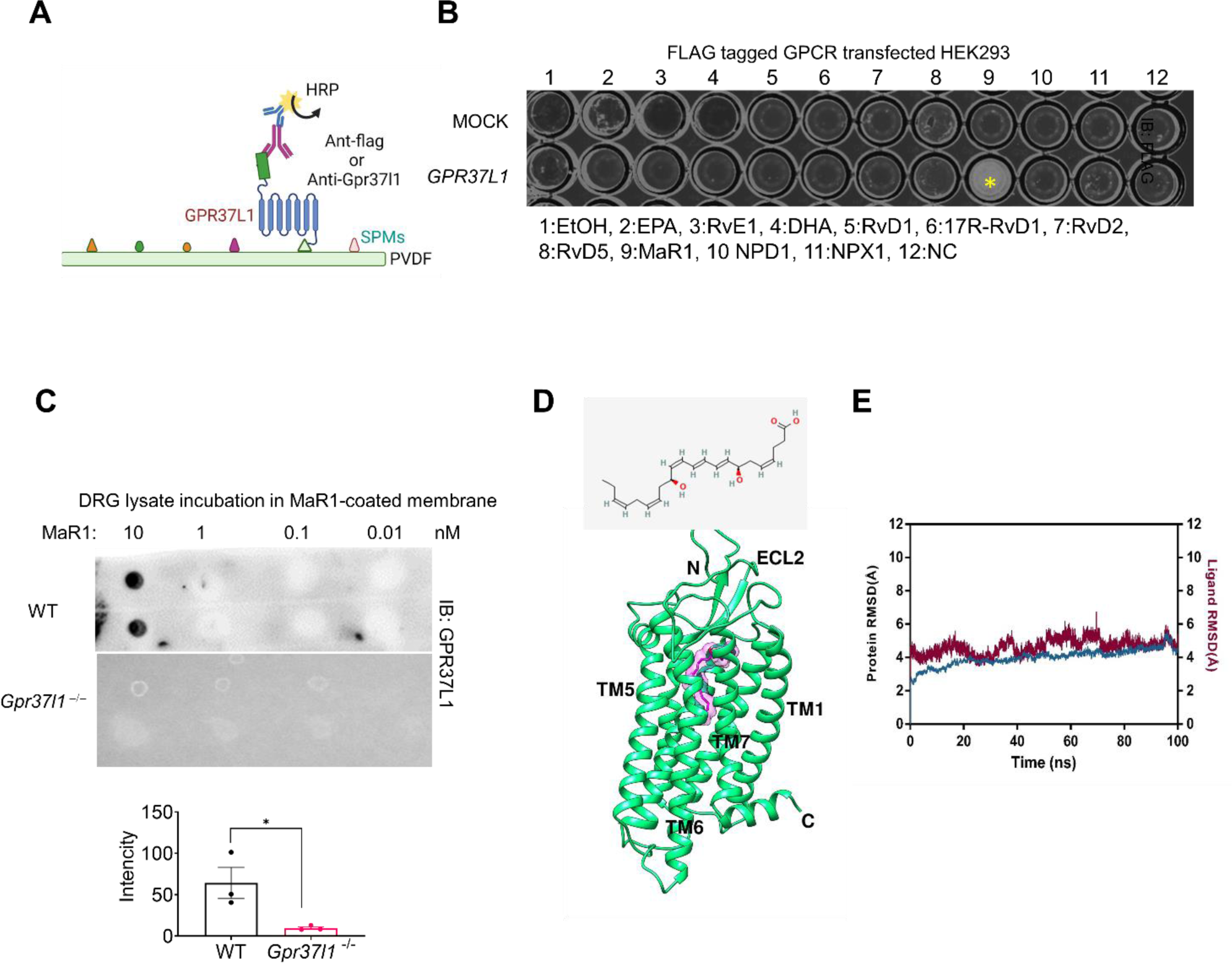
MaR1 binds GPR37L1. (**A, B**) Lipid overlay assay shows GPR37L1 binding to MaR1. (**A**) Schematic for detecting GPR37L1-binding lipid mediators. hGPR37L1 is FLAG-tagged and expressed in HEK293 cells after *hGPR37L1* cDNA transfection. (**B**) Specific binding of hGPR37L1 to MaR1. The plate was coated with 10 different lipid mediators (1 μg/ml, #2-11), as well as vehicle control (0.1% ethanol, #1) and no coating control (NC, #12), and then incubated with cell lysates of *hGPR37L1* or MOCK transfected cells. A yellow star indicates a positive response. **(C)** Up, a representative blot of MaR1-coated PVDF membrane. Down, quantification of dot intensity in DRG lysates from WT and *Gpr37l1* ^-/-^ mice. *n =* 3 repeats. **(D, E)** The overall structure of hGPR37L1 (Green) in complex with MaR1 (Magenta). **(E)** 100 ns RMSD graph was obtained with a 1000 ns simulation of the GPR37L1-MaR1 complex (red) or GPR37L1(blue). Data are expressed as mean ± s.e.m. and analyzed by unpaired t-test (C) \**P*<0.05.

Using GPR37L1 homology modeling, we generated computational predictions to examine the potential interaction between MaR1 and GPR37L1. The molecular structures for GPR37L1 and GPR37 have not been solved, but these two GPCRs have high homologies with human endothelial receptor Type B (human EDNRB, Supplemental Figure 5A) (31). Thus, we conducted EDNRB transmembrane helix sequence alignment with human GPR37L1 and utilized the crystal structure of Human EDNRB (PDB: 6IGK) as a template for GPR37L1 structural modeling (Figure 4D). We investigated possible interactions of MaR1 and NPD1 (control) with GPR37L1 by assessing the strength of hydrogen bonds between them using molecular docking and molecular dynamic simulation (MDS, Figure 4E, Supplemental Figure 5, B, and C). In the MDS of the GPR37L1-MaR1 complex cluster 1, the RMSD value of the protein backbone was stabilized at 4Å and the RMSD of MaR1 was between 4 to 6Å (Supplemental figure 5C). Further 100 ns simulation of the GPR37L1-MaR1 cluster-1 ensemble structure also demonstrated stable interactions, with RMSD values between 4 to 6Å (Figure 4D). In contrast, NPD1 does not show stable MDS activity for GPR37L1 (Supplemental Figure 5, B, and D), even though NPD1 can bind GPR37 (31).

### MaR1 inhibits STZ and PTX-induced pain via SGC and GPR37L1 signaling

MaR1 was shown to inhibit inflammatory pain in acute and chronic pain models (24, 26). We also tested the analgesic actions of MaR1 on STZ and PTX-induced neuropathic pain in *Gpr37l1*^+/+^, *Gpr37l1*^+/-^, and *Gpr37l1*^-/-^ mice. First, we administrated MaR1 via an intrathecal route that can target cells in DRGs and the spinal cord. We observed that, in *Gpr37l1*^+/+^ (WT) mice, MaR1 significantly reversed STZ-induced mechanical allodynia in all the mice (Figure 5A left, *P*<0.001). MaR1 also significantly reversed the mechanical allodynia in *Gpr37l1*^+/-^ mice (Figure 5A middle, *P* <0.05). However, MaR1 had no significant analgesic effects in *Gpr37l1*^-/-^ mice (Figure 5A right). Furthermore, we observed similar analgesic effects of MaR1 in the chemotherapy model (Figure 5B). MaR1 significantly reversed PTX-induced mechanical allodynia in WT mice (Figure 5B left, *P*<0.0001, *n =* 5). MaR1 also significantly reversed the mechanical allodynia in *Gpr37l1*^+/-^ mice (Figure 5B middle, *P* <0.01), but had no significant analgesic effects in *Gpr37l1*^-/-^ mice (Figure 5B right).

**Figure 5.**
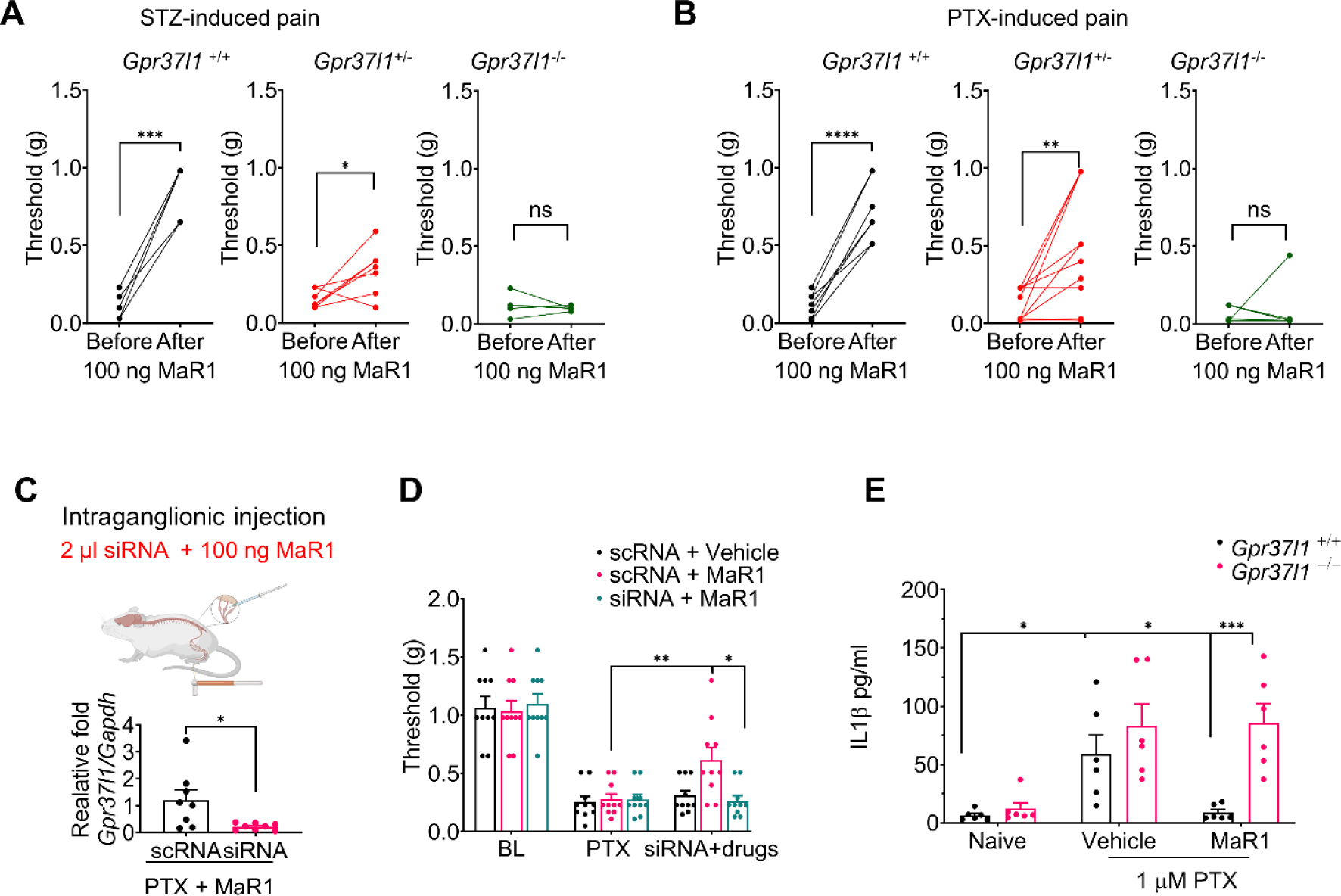
Intrathecal and intra-ganglionic injection of MaR1 reduces neuropathic pain via GPR37L1 expressed on SGCs. (**A, B**) Intrathecal MaR1 (100 ng) reduces mechanical allodynia in *Gpr37l1*^+/+^ and *Gpr37l1*^+/-^ mice, induced by 75 mg/kg of STZ (A) and 6 mg/kg of PTX (B). (**A**) Paw withdrawal thresholds were assessed in *Gpr37l1*^+/+^ mice (left, *n =* 7), *Gpr37l1*^+/-^ mice (middle, *n =* 10), and *Gpr37l1*^-/-^ mice (*n =* 8) after intrathecal MaR1 injection. (**B**) Paw withdrawal thresholds in *Gpr37l1*^+/+^ mice (left, *n =* 5), *Gpr37l1*^+/-^ mice (middle, *n =* 7), and *Gpr37l1*^-/-^ mice (*n =* 6) after intrathecal MaR1 injection. Behavior was assessed 1 h after MaR1 injection on Post-PTX and Post-STZ day 3. Abbreviations: PTX, paclitaxel; STZ, streptozotocin. (**C-D**) Validation of knockdown effect in L4 DRGs after scRNA or siRNA treated in PTX + MaR1 application (*n* = 10) **(C)**. I.G. injection of MaR1 (10 ng, 2 μl) reduces PTX-induced mechanical allodynia in control (scRNA-treated) animals but not in animals treated with *Gpr37l1*-siRNA **(D). (E)** MaR1 inhibits paclitaxel-induced IL-1β release in SGC-neuron co-cultures via GPR37L1. IL-1β release in neuron-glia cultures from DRG of WT and *Gpr37l1*^-/-^ mice was analyzed by ELISA. The cultures were stimulated with 1 µM paclitaxel for 24 h in the absence or presence of MaR1 (100 nM). *n =* 6 mice for WT and *n =* 6 mice for *Gpr37l1*^-/-^ mice. Data are expressed as mean ± SEM and statistically analyzed by paired t-test (A, B), Two-Way ANOVA with Bonferroni’s post-hoc test (D, E), Tukey’s post-hoc test (D, E) or unpaired t-test (C). \**P*<0.05, \*\**P*<0.01, \*\*\**P*<0.001, **** *P*< 0.0001, n.s., not significant.

Intrathecal injection of MaR1 could reduce pain at both DRG and spinal cord levels. To determine a direct effect of MaR1 on DRG cells, we conducted Intra-ganglionic (I.G) injection of MaR1 and *Gpr37l1* siRNA (Figure 5C). As expected, I.G. treatment of *Gpr37l1* siRNA resulted in a significant reduction of *Gpr37l1* mRNA expression (*P*<0.05, vs. scRNA, Figure 5C). Notably, I.G. injection of MaR1 significantly reduced mechanical pain in chemotherapy mice treated with control siRNA (scRNA, *P*<0.01), but this analgesic effect was compromised in mice treated with *Gpr37l1*-siRNA (*P*<0.05, Figure 5D). Collectively, these results indicate that MaR1 may relieve pain through GPR37L1 signaling in DRG.

SGCs have been shown to promote pain by releasing pro-inflammatory cytokines, such as IL-1β (8, 32). To examine the SGC-mediated neuro-glial interactions in a pathological condition, we prepared neuron-glia mixed cultures from the DRGs both WT and *Gpr37l1*^-/-^ mice and stimulated these co-cultures with 1 µM paclitaxel for 24 h. ELISA showed that PTX induced a marked increase in IL-1β levels in the culture medium and that this increase was blocked by 100 nM MaR1 in cultures from WT but not *Gpr37l1*^-/-^ mice (Figure 5E). Thus, MaR1 may partially alleviate neuropathic pain by suppressing IL-1β release via GPR37L1-mediated intracellular signaling in SGCs.

### MaR1 regulates KCNJ10 expression and K^+^ channel function in SGCs

KCNJ10 is the predominant potassium channel in mouse SGCs (3) (Supplemental Figure 1A) and is responsible for the generation of K^+^ currents in SGCs (33). Dysregulation of KCNJ10 in SGC has been implicated in the pathogenesis of pain (2). To define the direct effect of chemotherapy on the DRG, we employed an ex vivo whole-mount DRG preparation for both biochemical and electrophysiological analyses (Figure 6A) (34). We found that PTX treatment (1 μM) caused a rapid change in PM expression of expression (Figure 6B, Supplemental Figure 6, A and B) with significant down-regulation at 2 h (*P*<0.05, Figure 6, B and C, Supplemental Figure 6, A and B). Notably, 100 ng/ml MaR1 treatment was able to dramatically reverse this downregulation (*P*<0.05, Figure 6, B and C).

**Figure 6.**
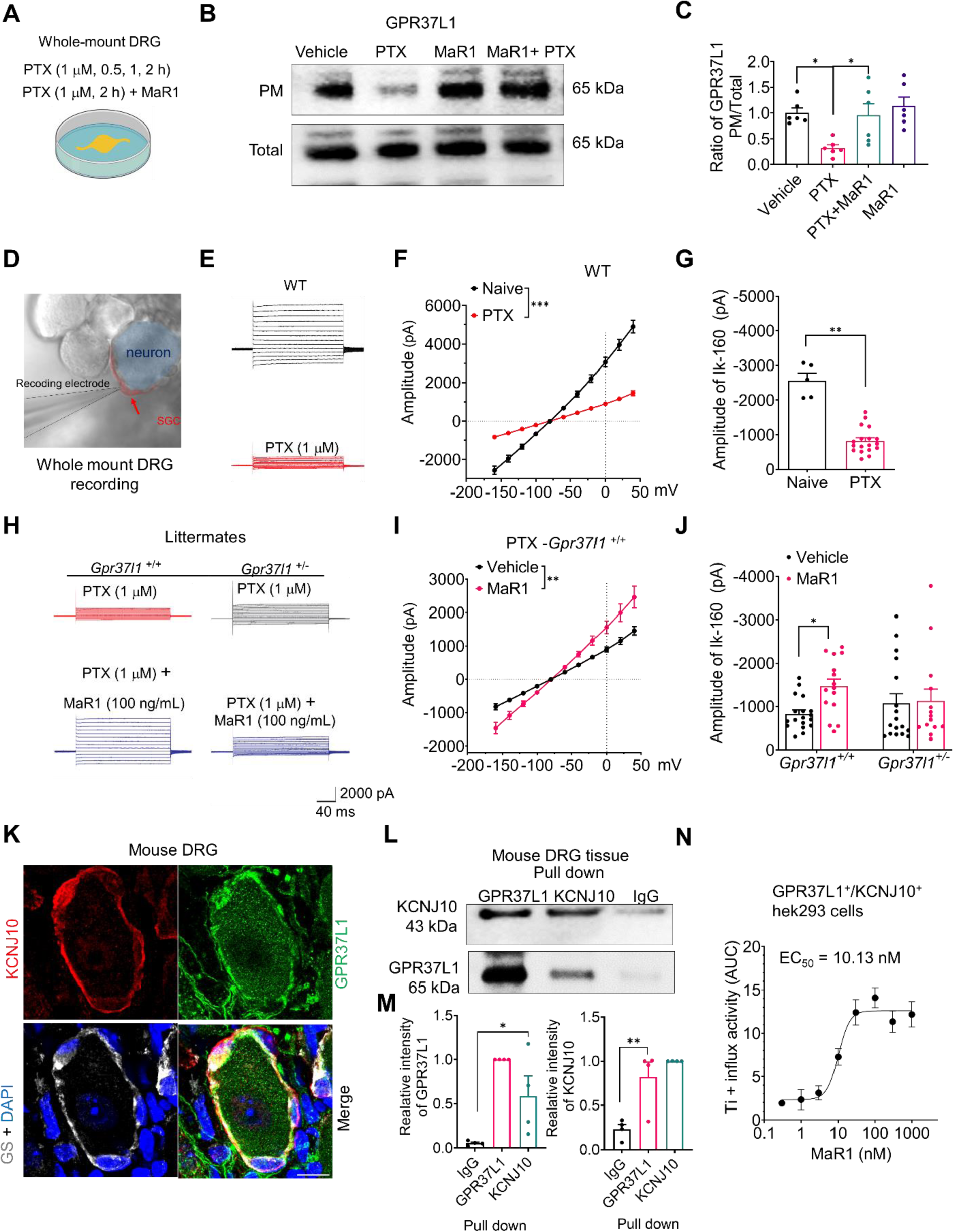
MaR1 increases the surface expression of GPR37L1 and KCNJ10/Kir4.1 and regulates K^+^ currents in SGCs via GPR37L1. **(A-C)** Western blot analysis of surface GPR37L1 expression in whole-mount DRG preparations. **(A)** Schematic of whole-mount DRG treatment with PTX or MaR1, followed by plasma membrane (PM) preparation and western blot. **(B**) Western blot showing the effects of PTX (1 μM) and MaR1 (100 ng/ml, 2 h) on PM and total fraction of GPR37L1. (**C**) Quantification of GRP37L1 expression (*n* = 6 preps). (**D-J**) Patch-clamp recordings of K^+^ currents in SGCs in whole-mount DRG preparations. (**D**) Micrograph showing patch-clamp recording in a SGC, indicated by a red arrow. (**E**) Representative trace for total K^+^ currents in SGCs of WT-DRG treated with vehicle or PTX (1 μM, 1 h). **(F)** Average I/V curves in SGCs treated with vehicle (*n* = 5 cells) and PTX (1 µM, *n* = 18 cells). **(G)** Quantification of the amplitude of K^+^ currents (I_k-160_) for F, with the holding potential of -160 mV. **(H)** Representative traces for total K^+^ currents in SGCs of *Gpr37l1*^+/+^ DRG treated with PTX (1 μM, 1 h) or PTX + MaR1 (100 ng/ml, 1 h), or in *Gpr37l1*^+/-^ DRG treated with PTX (1 μM, 1 h) or PTX + MaR1 (100 ng/ml). **(I)** Average I/V curves in WT SGCs after treatment of PTX (*n* = 18 cells) or PTX + MaR1 (*n* = 15 cells). **(J)** The amplitude of κ^+^ currents (I_k-160_) for H-I, with the holding potential of - 160 mV. In *Gpr37l1*^+/-^ DRG preps, *n* = 17 for PTX; *n* = 14 for PTX + MaR1. Note that the κ^+^ currents are suppressed by PTX and MaR1 can increase the suppressed currents in WT mice but not in mutant mice. **(K)** Triple IHC staining showing heavy co-localization of GPR37L1 (green), GS (White), and KCNJ10 (red) in mouse DRG SGCs encircling a neuron. Scale bars, 25 µm. **(L)** Protein pull-down and co-immunoprecipitation (Co-IP) showing GPR37L1/KCNJ10 interaction in mouse DRGs. Upper labels, pull-down antibodies; left labels, detection antibodies. **(M)** Quantification of the pull-down results in L for GPR37L1 (left) and KCNJ10 (right). *n* = 5 mice. **(N)** Dose-response curve of Ti^+^ influx after subtraction of background, as indicated by AUC (10 min, *n* = 6). Data are expressed as mean ± SEM and analyzed by Two-Way ANOVA with Bonferroni’s post-hoc test (F, I, and J) and One-Way ANOVA with Tukey’s post-hoc test (C, M), or two-tailed t-test (G). \**P*<0.05, \*\**P*<0.01, \*\*\**, P*<0.001. The EC_50_ was calculated by Richard’s five-parameter dose-response curve (N).

To investigate K^+^ signaling in SGCs, we conducted whole-cell patch-clamp recordings of K^+^ currents in SGCs (Figure 6, D-J). We visualized SGCs with DIC contrast microscopy (Figure 6D). Under normal conditions, SGCs exhibited large K^+^ currents (>2 nA) at the holding potential of -160 mV (*I*_k-160_, Figure 6E, Supplemental Figure 7, A and B). PTX treatment also caused a significant down-regulation of the surface expression of KCNJ10 at 2 h (*P*<0.001, Supplemental Figure 6, A and B).

To determine an acute effect of PTX on DRG cells and mechanical pain, we performed two local injections to target DRG cells via intra-ganglionic (I.G.) injection in the L4 and L5 DRG or intrathecal injection (Supplemental Figure 6, C-F). We measured acute mechanical pain at 1h and 2h after the local I.G. and I.T. injection. First, we found that I.G. injection of PTX was sufficient to induce mechanical allodynia within 1h, and furthermore, I.G. injection of 100 ng MaR1 could prevent the PTX-evoked acute mechanical pain at 1h (Supplemental Figure 6, C and D). Second, I.T. injection of PTX also evoked rapid mechanical allodynia within 1h, and PTX evoked mechanical allodynia was prevented by 100 ng MaR1 application in WT mice. Notably, MaR1 failed to inhibit I.T. PTX induced mechanical pain in Gpr37l1 knockout mice (Supplemental Figure 6, E and F). Collectively, these results demonstrated that 1) PTX is able to evoke acute mechanical pain via I.G. or I.T. injection and 2) MaR1 can prevent the PTX-induced acute pain via GPR37L1.

Strikingly, PTX (1 μM) induced a rapid reduction in K^+^ currents within 20 min (Figure 6, E-G; Supplemental Figure 7, B-D). PTX evoked a dose-dependent effect: a significant reduction of *I*_k-160_ was observed at concentrations as low as 0.1 μM PTX (*P*<0.05, Supplemental Figure 7E). As previously reported (35), the total K^+^ currents were almost completely blocked by the Kir channel blocker Barium (100 μM); the amplitude of the Kir-mediated *I*_k-160_ current was comparable with that of the total K^+^currents (Supplemental Figure 7, F and G). Co-application of MaR1 (100 ng/ml) with PTX significantly reversed the K^+^ current deficit (*P*<0.01, Figure. 6, H and I). Importantly, MaR1’s enhancement of K^+^ currents was abolished in SGCs of *Gpr37l1* mutant mice (Figure. 6, H and J; Supplemental Figure 7H).

To investigate how GPR37L1 regulates KCNJ10 mediated K^+^ channel function, we performed: 1) IHC to confirm GPR37L1/KCNJ10 co-expression in mouse DRG sections (Figure 6K), 2) GPR37L1/ KCNJ10 Co-IP in mouse DRG tissues (Figure 6, L and M), and 3) Ti^+^ influx assay for assessing intracellular K^+^ levels in HEK293 cells transiently expressing GPR37L1 and KCNJ10 (Figure 6N, Supplemental Figure 7I). Triple IHC staining revealed that KCNJ10 is highly co-expressed with GPR37L1 in GS^+^ SGCs (Figure 6K). Co-IP analysis revealed that GRP37L1 or KCNJ10 antibodies could pull down KCNJ10 or GPR37L1 respectively in mouse DRG lysates (Figure 6, L and M). Ti^+^ influx assay showed a dose-dependent increase in Ti^+^ influx in HEK293 cells incubated with 0.3 to 1000 nM (1 h) of MaR1, with an EC_50_ = 7.2 nM (Figure 6N). MaR1-induced Ti^+^ influx peaked in 10 min (Supplemental Figure 7J). The acute paclitaxel evoked mechanical allodia also can block the 100 ng Mar1 application (Supplemental Figure 7K n = 6). Collectively, these results demonstrate that MaR1 regulates KCNJ10-mediated K^+^ influx through GPR37L1 signaling.

### MaR1 regulates KCNJ3 expression and K^+^ channel function in human SGCs

We used a published database of scRNA seq of mouse and human TG (36, 37) to compare the species differences in expression levels and cell types of the genes encoding KCNJ family of K^+^ channels in SGCs of mouse TGs (Figure 7A) and human TGs (Figure 7B). In mouse TGs, *Kcnj10 i*s predominantly expressed by SGCs, although Schwann cells and immune cells show moderate levels of *Kcnj10* expression (Figure 7A). Compared to the mouse gene, the human *KCNJ10* expression is lower, but still specific for SGCs (Figure 7B). We found a striking species difference in KCNJ3 expression in mouse and human TGs. *Kcnj3* is expressed by sensory neurons but not by SGCs in mouse TGs (Figure 7A). In sharp contrast, *KCNJ3* is highly expressed by SGCs, with some low expression in sensory neurons and Schwann cells in human TGs (Figure 7B). Cluster analysis (36) showed that *Gpr37l1* is associated with *Kcnj10* in mouse TGs and *GPR37L1* is associated with both *KCNJ3* and *KCNJ10* in human TGs (Figure 7, A and B). Additional analysis of mRNA expression of human DRGs of DPN and control patients from a published database (29) revealed significant downregulations in the mRNA expression levels of *GPR37L1* (*P*<0.01) and *KCNJ3* (*P*<0.05), but not *KCNJ10*, in the DPN group (Figure 8C). ISH analysis showed that both *KCNJ3* and *KCNJ10* are expressed by SGCs, surrounding *TUBB3*^+^ neurons, in human DRG sections (Figure 8D). Further analysis of human RNA-seq data (36) revealed a higher percentage of colocalization for *GPR37L1*/*KCNJ3* (∼75%) than *GPR37L1*/*KCNJ10* (∼30%) (Supplemental Figure 8A). Western blot analysis showed a significant decrease of KCNJ3 in the plasma membrane of DRG samples from neuropathic pain patients (*P*<0.01, vs. Control, Figure 7, E and F).

**Figure 7.**
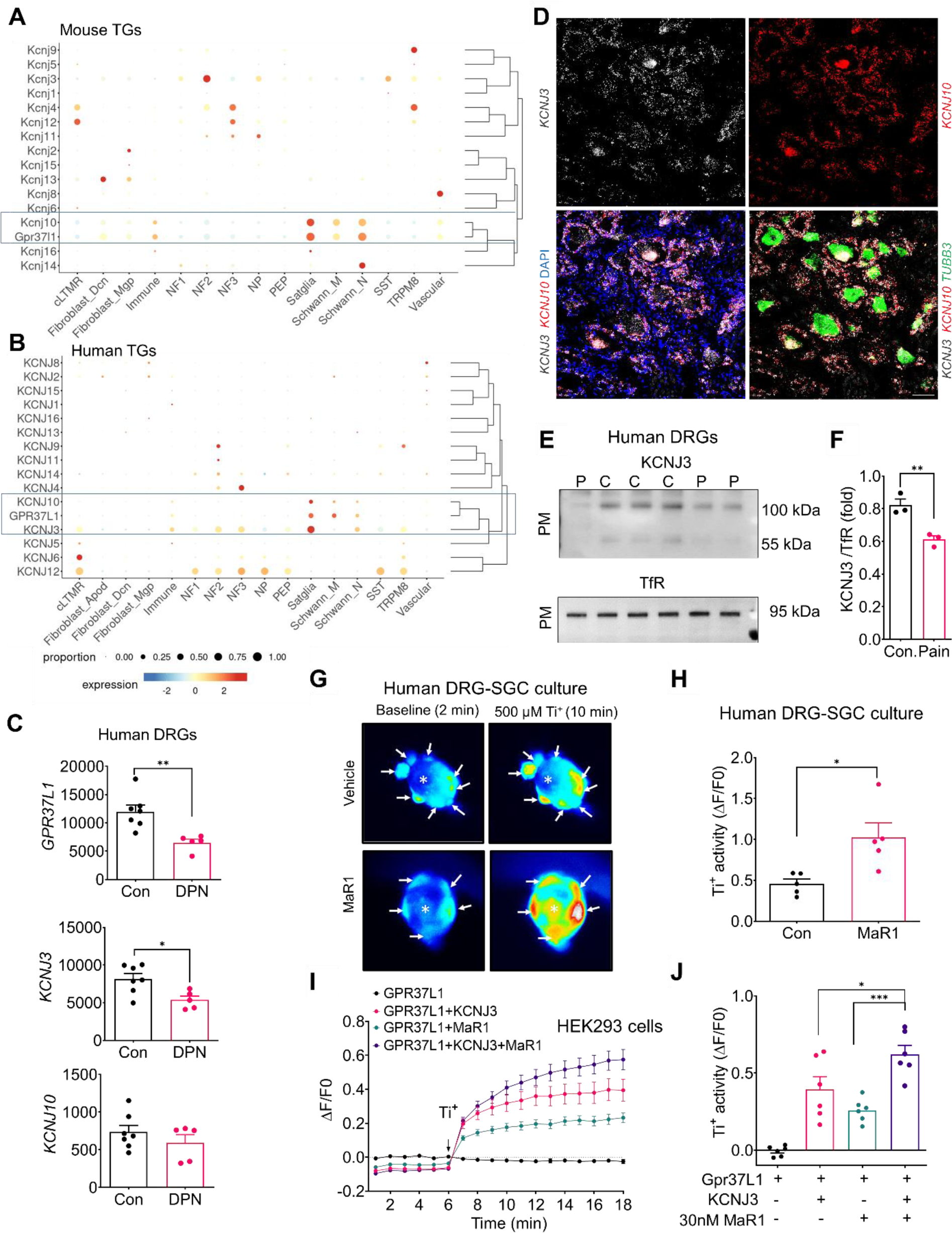
GPR37L1 regulates KCNJ3/GIRK1 signaling in human SGCs. **(A, B)** Single-cell RNAseq meta-analysis of KCNJ transcripts expression in mouse (A) and human (B) trigeminal ganglia (TGs) from a published database (36). (**A)** Mouse SGCs predominantly express *Gpr37l1* and *Kcnj10*, which are clustered together. **(B)** Human SGCs express *GPR37L1, KCNJ3*, and *KCNJ10*, which are clustered together. Note greater expression *KCNJ3* than *KCNJ10*. **(C)** Transcriptomic meta-analysis of human DRGs, from a published database (29), reveals downregulations of *GPR37L1* and *KCNJ3*, but not *KCNJ10*, in patients with painful DPN. **(D)** Triple RNAscope in situ hybridization (ISH) showing the expression of *KCNJ10* (red), *KCNJ3* mRNA (white), and *TUBB3* (Green) in non-diseased human DRG. Note co-localization of *KCNJ10* and *KCNJ3* in SGCs surrounding the *TUBB3*+ neurons. Scale bar, 50 µm. **(E)** Western blots showing PM fractions of KCNJ3 levels in human DRGs of neuropathic pain patients and controls. **(F)** Quantification of the western blot results in E (*n* = 4). (**G, H**) MaR1 increases K^+^ levels in human SGCs. **(G)** Images of Ti^+^ influx in human DRG co-cultures for SGCs and neurons treated with vehicle and MaR1. SGCs are indicated with white arrows. **(H)** Quantification of fluorescence intensity for G at 10 min after 500 µM Ti^+^ simulation in human SGCs. *n* = 5 cultures from 2 donors. **(I)** Time course of Ti^+^ influx in HEK293 cells expressing GPR37L1 with vehicle (*n* = 16) or MaR1(100 nM, 30 min *n* = 16) and GPR37L1/KCNJ3 in the presence of vehicle(*n* = 13) and MaR1 (100 nM, 30 min, *n* = 12). **(J)** Quantification of Ti^+^ influx in I after 10 min of 500 µM Ti^+^ stimulation (*n* = 6 culture). Data are expressed as mean ± SEM and analyzed by two-tailed t-test (C, F, H) or One-Way ANOVA with Tukey’s post-hoc test (J). \**P*<0.05, \*\**P*<0.01, \*\*\**P*<0.001.

**Figure 8.**
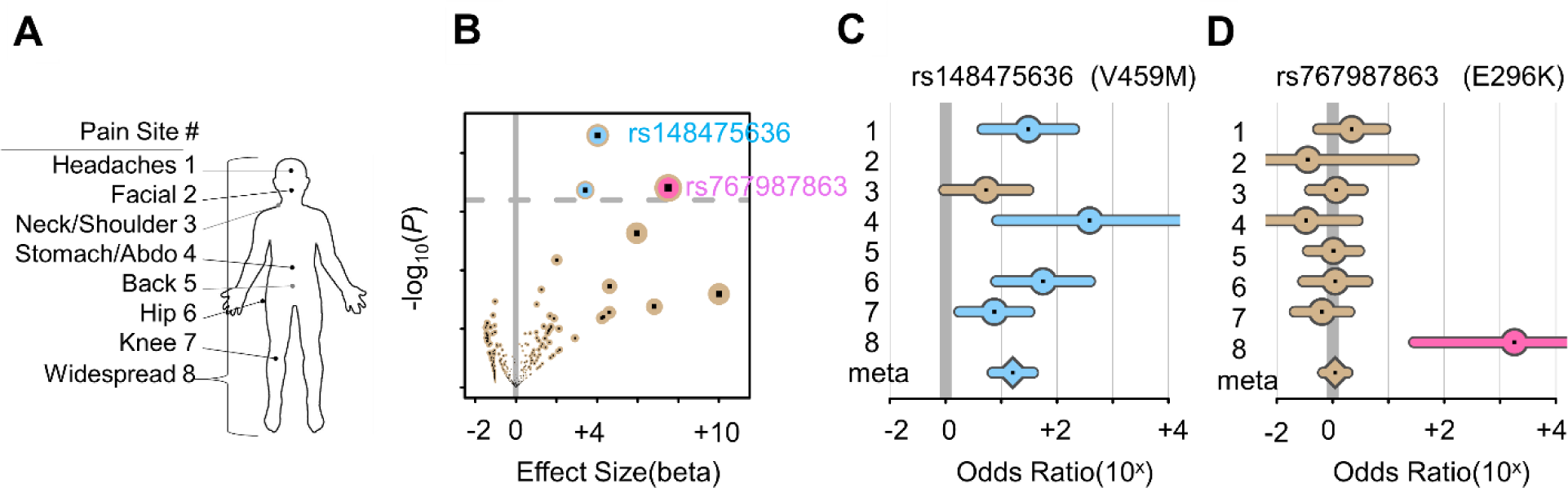
*GPR37L1* rare variants effects in chronic pain. (**A**) The eight chronic pain sites found in the UK Biobank project. (**B**) Volcano plot showing rare variants’ significance as a function of effect size. Each dot is a variant. The vertical bar indicates the null effect, while the dotted horizontal bar indicates the threshold for a false discovery rate of 20%. Two significant variants were identified: rs148475636 (blue) and rs767987863 (pink). (**C**) Forest plot for variant rs148475636. **(D)** Forest plot for variant rs767987863. The forest plots show variants’ effects at the eight chronic pain sites. Segments track a 95% confidence interval for point estimates of odds ratios. Missing estimates when allele counts less than five. Meta estimate from a meta-analysis of all available pain sites. Segments are colored (blue/pink) when *P* <0.05

KCNJ3 K^+^ channel is known to be modulated by GPCRs, and the Ti^+^ influx assay has been used for G protein (βγ subunit-mediated) drug screening (38). To investigate whether GPR37L1 regulates K^+^ influx via KCNJ3, we conducted a Ti^+^ influx assay in both human SGCs (Figure 7, G and H) and HEK293 cells expressing GPR37L1 or GPR37L1 together with KCNJ3 (Figure 7I). Human DRG culture imaging revealed an increased dye signal in SGCs surrounding neurons following 0.5 mM Ti^+^ incubation. However, MaR1 treatment (100 nM, 0.5 h) caused significant increases in K^+^ signaling in human SGCs (*P*<0.05, vs. Control, Figure 7, G and H; Supplemental Figure 8B). In GPR37L1/KCNJ3-expressing HEK293 cells, MaR1 treatment was sufficient to increase K^+^ levels (Figure 7I). As expected, KCNJ3 over-expression was also able to increase K^+^ levels in GPR37L1-expressed cells. However, MaR1 caused a further increase in K^+^ levels in KCNJ3/GPR37L1-expressing cells (Figure 7, I and J). Finally, MaR1-induced K^+^ increase in KCNJ3/GPR37L1-expressing cells was blocked by either 1 µM of KCNJ3 antagonist SCH 23390 (39) or 1 µM of G_βγ_ inhibitor (Gallein) (40) (Supplemental Figure 8, C and D). MaR1 alone had no effect on Ti^+^ influx in KCNJ3-expressing cells. In contrast, the positive control ML-297, a KCNJ3 agonist (300 nM), was sufficient to induce Ti^+^ influx in KCNJ3-expressing cells (Supplementary Figure 8, E and F).

### *GPR37L1* mutations are associated with chronic pain in humans

We examined the genetic correlation of *GPR37L1* variants with chronic pain. We assessed the UK Biobank cohort to inquire about the potential roles of rare GPR37L1 variants in chronic pain, as it offered accurate allele assignments via whole-exome sequencing (41). SAIGE was used to perform association tests between rare coding variants and a report of chronic pain at eight body sites (Figure 8A and Supplemental Table 3A). The rare coding variants were chosen for analyses because they are most likely to have a significant and interpretable effect and they usually result in a reduction of the function of the corresponding protein rather than an increase. Furthermore, they provide a precise target for follow-up functional analysis (42). A total of 380 variant-site associations with at least five minor allele counts were considered in a primary analysis (Figure 8B). Of these, three were found significant at the false discovery rate level of 20%: rs148475636 (V459M) was associated with hip pain and headache (OR=56.1, *P* = 5.0×10-5, and OR=30.5, *P* = 3.95×10-4) and rs767987863 (E296K) was associated with widespread pain (OR=1816, *P* = 3.9×10^-4^) (Supplemental Table 3A). We then performed secondary analyses to estimate the risk for chronic pain at all eight sites for two identified variants (Figure 8, A-D). The variant of SNP rs148475636 was associated with significantly increased risk at many body sites, suggesting a pleiotropic effect, including an overall significant and large risk in a meta-analysis (meta OR=16, *P*=1.0×10-10), while a meta-analysis on SNP rs767987863 was not significant (Figure 8, A and D, Supplemental Table 3B). Importantly, both significant SNPs are non-synonymous and display high CADD scores (43), suggesting that the variants alleles are deleterious and have a loss of function (8.0 for rs148475636 and 27.3 for rs767987863; Supplemental Table 3A). More broadly, the majority of the tested coding variants had substantial pathogenicity CADD scores (Supplemental Table 3A) and were risk factors for chronic pain (Figure 8D), suggesting that GPR37L1 is protective against pain. Further, in-slico prediction of protein stability revealed that among the six mutations of *GPR37L1* variants, the allele E296K showed a significant loss of protein stability (*n* = 9 simulation, Supplemental Figure 9A).

We further characterized the effects of the E296K mutation on GPR37L1 and KCNJ3 expression, IL-1β release, and Ti^+^ influx in human SGC cultures (Supplemental Figure 9, B-D). In E296K-expressed HEK cells, we did not observe significant changes in GPR37L1 and KCNJ3 expression (E296K vs. WT, Supplemental Figure 9B). However, E296K mutation resulted in increased IL-1β release (WT vs. E296K) following PTX treatment (1 μg/ml, 12 hours, Supplemental Figure 9C). Furthermore, MaR1-induced Ti^+^ influx activity was abolished in E296 mutant-expressing human SGCs (*P*<0.01, compared with WT, Supplemental Figure 9D). Collectively, these findings suggest that the E296K mutation of *GPR37L1* may decrease protein stability, increase IL-1β production, and abrogate MaR1-mediated regulation of K^+^ channel function, leading to increased risk of developing chronic pain.

## Discussion

In this study, we have offered several new insights into GPR37L1 and satellite glial signaling in pain control. First, we demonstrated a protective role of GPR37L1 against chronic pain, as neuropathic pain caused by chemotherapy or STZ-induced pain is enhanced and prolonged in transgenic mice with *Gpr37l1* deficiency (*Gpr37l1*^+/-^ and *Gpr37l1*^-/-^). We also showed that partial knockdown of GPR37L1 in DRGs of naïve wild-type animals was sufficient to induce mechanical allodynia. Furthermore, we found GPR37L1 downregulations not only in DRGs of mice with chemotherapy and diabetes but also in DRGs of patients with painful diabetic neuropathy. Importantly, we have identified novel SNPs of human *GPR37L1* and potential variant alleles that are deleterious with loss of function, supporting our functional studies in mice that GPR37L1 is protective against the development of chronic pain.

Another important finding is that we identified MaR1 as a novel ligand of GPR37L1. Prosaposin, a highly conserved glycoprotein and secretory protein, was initially implicated as a ligand for GPR37 and GPR37L1 (10). Single-cell based pathway analysis in mouse DRGs also revealed an association of prosaposin and *Gpr37l1* (44). However, it is unclear how prosaposin interacts with GPR37L1. Our data showed that *Gpr37l1* and *GPR37L1* transcripts are among the top 10 expressed GPCR transcripts in mouse and human DRGs. Given its high expression under the physiological conditions, constitute activity of GPR37L1 was proposed, and metalloprotease cleavage of the N terminal of GPR37L1 may reduce its constitutive activity (12). It was also suggested that this orphan receptor remains unliganded (13). MaR1 is a DHA-derived SPM that has exhibited potent analgesic actions in animal models of inflammatory pain (24), arthritic pain (26), and postoperative pain (45). So far, all the known SPM receptors are GPCRs and each SPM may have multiple receptors (23). It was found that the GPCR LRG6 acts as a specific receptor for MaR1, and activation of LRG6 by MaR1 promoted phagocyte immune resolvent functions (46). Notably, LRG6 was identified by β-arrestin assay, and GPR37L1 showed no such activity (46). Our data showed a novel signaling of MaR1 via GPR37L1. The lipid overlay and protein pulldown experiments demonstrated MaR1 binding to GPR37L1. Furthermore, the computer simulations revealed that MaR1 forms hydrogen bonding with ASN190, ARG196, and GLU375 residues in the GPR37L1 binding pocket, whereas NPD1 only showed weak binding to this receptor. Importantly, our functional evaluations demonstrated that GPR37L1 is required for MaR1’s analgesic actions in mouse models of neuropathic pain.

Our results demonstrated that GPR37L1 controls neuropathic pain in mice by regulating the surface expression and function of KCNJ10 (Kir4.1) in SGCs. We have provided a new mechanism by which chemotherapy and diabetes induce pain via SGC signaling. Chemotherapy caused a rapid suppression of KCNJ10-mediated K^+^ currents in SGCs. Remarkably, MaR1 conferred protection against CIPN by inducing GPR37L1-dependent upregulations of surface KCNJ10 expression and KCNJ10-mediated K^+^ currents in SGCs. We postulate that dysregulation of GPR37L1/KCNJ10 signaling in SGCs in neuropathy conditions (CIPN and DPN) will impair the SGC’s function of K^+^ buffering, leading to sequential increases in extracellular levels of K^+^, nociceptor excitability, and pain sensitivity. Importantly, this dysregulation can be prevented and treated by GPR37L1 agonists such as MaR1 (Supplemental Figure 10). Additionally, MaR1 also blocked paclitaxel-induced IL-1β release in SGC-neuron co-cultures. Control of neuroinflammation in DRG by the MaR1/GPR37L1 axis will further reduce neuropathic pain.

Development of pain medicine has been hampered by the translational gap between rodents and humans, as many genes are differentially expressed and functioned in mouse and human tissues including DRGs (47-50). We validated critical role of the MaR1/GPR37L1 axis in both mouse (SGC) and human cells (SGC and HEK cells). Both *Gpr37l1* and *GPR37L1* mRNAs are highly expressed in mouse and human DRGs. We also found striking species differences in KCNJ3 and KCNJ10 expression in mouse and human TGs. In mouse sensory ganglia, *Kcnj3* is expressed by neurons but not by SGCs (Figure 7A) (17, 51). However, in human sensory ganglia, *KCNJ3* mRNA is highly enriched in SGCs (Figure 7B). This distinction highlighted different mechanisms of KCNJ3 in pain modulation in mice and humans. As an inwardly-rectifying potassium channel (GIRK), KCNJ3 (GIRK1) regulates the G-protein mediated outward K^+^ currents in neurons induced by analgesics, such as opioids (52). Importantly, KCNJ3 is coupled to GPR37L1 in human SGCs and its activity is positively regulated by MaR1. Although KCNJ10 (Kir4.1) is expressed by SGCs of both mouse and human sensory ganglia, its contribution to K^+^ buffering could be greater in mouse than human. In mouse sensory ganglia GPR37L1 is only associated with KCNJ10. However, in human sensory ganglia, GPR37L1 is associated with both KCNJ10 and KCNJ3, with much higher co-localization with KCNJ3 than KCNJ10 (Supplemental Figure 8A). Despite these species’ differences, the MaR1/GPR37L1 pathways are critical for SGC K^+^ signaling in both mice and humans.

Neuropathic pain following DPN and CIPN poses a significant health problem (53-55). (56, 57). Current treatments for neuropathic pain are inadequate and often accompanied by substantial side effects. Our findings propose that directing therapeutic focus toward GPR37L1 in SGCs could develop novel approaches for addressing neuropathic pain post-CIPN and DPN. MaR1 is a highly potent GPR37L1 activator and has a wide safety profile, but its pharmacokinetics and in vivo stability (e.g., short half-life as a lipid mediator) remain to be improved. An alternative is to identify novel small-molecule agonists of GPR37L1 for pain management. Although we have shown specific and high expression of GPR37L1 in mouse and human SGCs, astrocytes in the central nervous system also express GPR37L1, which plays a protective role in astrocytes (10). The study of GPR37L1’s function and its activation by MaR1 in astrocytes is of great interest. Notably, SGCs and astrocytes reside in the PNS and CNS, respectively, and play similar and distinct roles in pain regulation (6). Given the well-known side effects of CNS-targeting drugs, targeting SGC-GPR37L1 in the PNS may offer a safer analgesic drug for pain relief and disease modification.

## Materials and Methods

### Reagents

Maresin 1 (MaR1, Cat No. 10878) and other lipid mediators were purchased from Cayman Chemical. *GPR37L1*-Tango plasmid was from Addgene (Cat No.66356). Human *GPR37L1-WT* and human *GPR37L1-E298G* mutant cDNAs were synthesized from Twist Bioscience (South San Francisco, CA, US). The human *KCNJ10-v5* plasmid was from DNASU (Clone: HsCD00939635). Human *KCNJ3-HA* plasmid was purchased from Applied Biological Materials (Richmond, BC, Canada; Cat no: 253840610295). *Gpr37l1* siRNA and control scRNA were purchased from Ambion life technology (Cat No. 173005 for *Gpr37l1*, Cat No, 4459405 for *scRNA*), and transfection agent PEI/RNA polyplexes (LNP102) were from ABP bioscience (Cat No. LP007-1). Flag-tagged *human GPR37L1* was purchased from Origene (Cat No. RC208132). *Fabp7-mGpr37-*AAV*9* or mock AAV9 virus was generated by Vector Builder (Chicago, IL, US). The Brilliant Thallium flex kit (Cat No. 11000-10) was purchased from Ion Bioscience (San Marcos, TX, US;). Paclitaxel (Cat No. T719) and streptozotocin (Cat No. S0130) were purchased from Sigma-Aldrich. SCH 23390 (Cat No. 0925), ML297 (Cat No. 5380), and Gallein (Cat No. 3090) were purchased from Tocris.

### Animals

*B6;129S5-Gpr37l1^tm1Lex^/Mmucd* strain was obtained from UC Davis (MMRRC, stock Cat No. 011709-UCD). *Gpr37l1* and littermate mice with C57BL/6 background were maintained at Duke University Medical Center. CD1 mice were also used for some behavioral, histochemical, and electrophysiological studies. Adult mice (males and females, 8–10 weeks) were used for behavioral tests and biochemical assays. Both sexes were included in behavioral testing and no sex differences were noticed in behavioral and cellular tests conducted in this study. Two to five mice were housed in each cage under a 12-hour alternating light-dark cycle with ad libitum access to food and water. Sample sizes were estimated based on our previous studies for similar types of behavioral and biochemical analyses (28). Animal experiments were conducted according to the National Institutes of Health Guide for the Care and Use of Laboratory Animals and approved by the Institutional Animal Care and Use Committee (IACUC).

### RNA purification, sequencing, and analysis

To perform poly-A RNA-sequencing, L1-L5 mouse DRG was homogenized with an RNeasy Mini Kit (Qiagen, Valencia, CA) using on-column DNase-I digestion according to the manufacturer’s protocol. RNA quality was assessed using a Nano-drop (Thermo Scientific) and RNA sequencing service was requested from LC science (Houston, TX, US). For the mouse DRG RNA-seq analysis, paired-end sequencing was performed with a read length of 150 bp. Read counts per gene per sample were quantified using HTSeq version 0.9.1. The detected total gene number is 25.091–25.664 in DRG samples.

GPCR mRNA expression levels in human DRGs were taken from the human DRG expression quantitative trait loci study (58). In short, DRG expression levels were determined from total RNA extracts measured using Affymetrix’ Human Transcriptome Array 2.0 (Affymetrix, Santa Clara), scanned with Affymetrix’ GeneChip 3000 G7 Instrument System, then probe intensities were normalized in a standard fashion.

### Pain models and drug injection

Mouse models of chemotherapy-induced peripheral neuropathy (CIPN) were induced by paclitaxel, given via intraperitoneal route either by a single injection (6 mg/ml, i.p.) or by 4 injections (2 mg/kg, i.p.) on days 0, 2, 4, 6 (59). A mouse model of diabetic peripheral neuropathy (DPN) was induced by streptozotocin (STZ) via an intraperitoneal injection (75 mg/kg) (60). For intrathecal (i.t.) injection of MaR1 or PTX, a spinal cord puncture was made by a Hamilton micro-syringe (Hamilton) with a 30-gauge needle between the L5 and L6 levels to deliver reagents (5 μl) to the cerebral spinal fluid (61).

### Intra-ganglionic injection of siRNA, PTX, MaR1, and AAV-virus

Mice were placed in a prone position under isoflurane anesthesia and a 1 cm posterior longitudinal skin incision was made in the lumbar portion of the spine. After the plate of the ipsilateral L5 vertebra was carefully removed to expose the L5 spinal nerve and the L4-L5 DRGs, microinjection of 1-2 µl of the solution was administrated to L4 and L5 DRG using a Hamilton syringe (80030) connected to a glass micropipette (tip diameter 10–20 μm) (62). After injection, the wound was covered with a gelatin sponge solution to prevent leakage and then closed with a suture. The following reagents were injected with gelatin spongey (2 µl): 1) 3 μg of *Gpr37l1*-targeting siRNA or scramble RNA, mixed with PEI/RNA polyplexes (1 µl LNP102); 2) MaR1 (100 ng) or PTX (100 ng); 3) AAV9-*Fabp7*-*mouse Gpr37L1* (1 x 10^12^).

### HEK293 cell culture and *GRP37L1, KCNJ10,* and *KCNJ3 t*ransfection

The HEK293 Flp-In^TM^ cell line (Invitrogen, R78007) was purchased from the Duke Cell Culture Facility. Cells were cultured in high glucose (4.5 g/L) Dulbecco’s Modified Eagle’s Medium containing 10% (v/v) fetal bovine serum (Gibco). The *GPR37L1*, *GPR37L1-WT*, *GPR37L1-E296K*, *Mock* control, *KCNJ10,* and *KCNJ3* cDNAs were transfected using Lonza 4D-Nucleofector^TM^ X-unit (2 µg cDNA/1 x 10^7^ cells; protocol No.CM130). After reaching 70% confluency by co-transfection with 1 µg of pmaxGFP control cDNA (Lonza), the transfected cells were cultured for an additional 48 h before use.

### SGC cultures and SGC-neuron co-culture

DRGs were collected from both WT and KO mice (8 weeks male) and were incubated in 1 mg/ml Collagenase/Dispase (Roche Diagnostics, Madison, WI, USA) at 37℃ for 60 min, with agitation at 100 RPM and then followed by incubation in 0.05% trypsin/EDTA for 10 min. The digestion enzymes were prepared in Dulbecco’s modified Eagle medium/F12 with GlutaMAX (ThermoFisher). After incubation with 0.1% trypsin inhibitor and centrifugation (300G), the cell pellet was gently triturated in a neurobasal medium containing 0.5 µM glutamine. Dissociated DRG cells were seeded on non-coated culture dishes for 4 h. SGCs and neurons were included for co-culture by hand-shaking of the culture flasks gently for 5 to 10 min and then resuspended by replacing them with a new neurobasal medium. The attached glial-like cells were cultured in the DMEM/F12 medium containing 10% fetal bovine serum and 1% streptomycin/penicillin to promote cell growth and inhibition of differentiation. After 2 to 6 days in culture, the glial-like cells were differentiated by application of the serum-free neural basal medium. The neuron-rich fraction was seeded on a Poly L lysine (Sigma) coated plate with a Neurobasal medium containing 2% B27 (ThermoFisher) (63).

### Lipids overlay assay

Lipids membrane coating and protein overlay assay were performed as previously described (64). Lipid mediators including SPMs (RvD1, RvD2, RvD3, NPD1, MaR1), DHA, and vehicle EtOH, were directly loaded on hydrophobic PVDF membrane walls (96 well plates; Bio-Rad). Compound-coated membranes were dried and blocked with 1% BSA. The coated membranes were incubated with lysates obtained from *hGPR37L1-*transfected HEK293 cells or with DRG lysates from WT or *Gpr37l1* KO mice for 2 h, followed by detection using an anti-GRP37L1 antibody (Bioss, Rabbit, 1:1000, Cat No. bs-15390R) or anti-flag antibody (Cell signaling, Rabbit, 1:1000, Cat No.14793). Blots were further incubated with an HRP-conjugated secondary antibody (Anti-rabbit, Jackson ImmunoResearch, raised in donkey, 1:5000), developed in ECL solution (Pierce), and protein signal was visualized from ChemiDoc XRS (BioRad). The signal intensity was quantified by Image J software (NIH).

### Lipid pull-down assay

Isolated membrane proteins were pre-adsorbed to uncoated control agarose beads (Vector labs); the unbound fraction was collected and incubated for 24 h at 4°C with lipid-coated agarose beads (28). After extensive washing, bound proteins were eluted from the lipid-coated beads, suspended in Laemmli buffer containing 2% SDS (w/v) and 0.3M β-mercaptoethanol (Sigma), and heated for 5 min at 95°C to dissolve proteins before separation on 4–20% polyacrylamide/SDS gels. The lysate proteins were detected by western blot.

### Co-immunoprecipitation

Mouse DRGs were harvested, washed once in ice-cold PBS, and then scraped in harvest buffer (10 mM HEPES, 100 mM NaCl, 5 mM EDTA, 1 mM benzamidine, protease inhibitor tablet, 1% Triton X-100, pH 7.4). Cell lysates were then solubilized, and immunoprecipitated with anti-GPR37L1 antibody (Bioss, Rabbit, 1:100, Cat No. bs-15390R), anti-Kir4.1/KCNJ10 antibody (Thermo Scientific, Rabbit, 1:100, Cat No. 12503-1-AP), or normal rabbit IgG (Santacruz,1:100, Cat No. sc-2027). The antibody-binding proteins were pulled down using protein A conjugated magnetic beads (Thermo Scientific, Cat No.88845) and washed by repeated centrifugation and homogenization. Samples were heated, then probed by Western blotting using anti-GPR37L1 (Alomone Labs, Rabbit, 1:1000, Cat No.AGR-050) or anti-Kir4.1 (Thermo scientific, Guinea pig, 1:1000, Cat No. PA5-111798).

### Human DRG samples

Non-diseased human DRGs were obtained from donors through the National Disease Research Interchange (NDRI) with permission of exemption from the Duke University Institutional Review Board (IRB). Human DRGs from neuropathic pain patients and healthy controls were obtained from a cohort collected at the University of Pittsburg (58). They consist of a snap-frozen bilateral lumbar L4 and L5, collected from brain-dead subjects following asystole with the consent of first-tier family members. All procedures were approved by the University of Pittsburgh Committee for Oversight of Research and Clinical Training Involving Decedents and the Center for Organ Recovery and Education, Pittsburgh, PA (http://www.core.org).

### Human SGC culture and SGC-neuron-rich culture

Human DRG cultures were prepared as previously reported (65). DRGs were digested at 37°C in a humidified CO_2_ incubator for 120 min with collagenase Type II (Worthington, 290 units/mg, 12 mg/ml final concentration) and Dispase II (Roche, 1 unit/mg, 20 mg/mL) in PBS with 10 mM HEPES, pH adjusted to 7.4 with NaOH. DRGs were mechanically dissociated using fire-polished pipettes, filtered through a 100-m nylon mesh, and centrifuged (500 g for 5 min). The pellet was resuspended, plated on 0.5 mg/mL poly-D-lysine–coated glass coverslips, and grown in Neurobasal medium supplemented with 10% FBS, 2% B-27 supplement, and 1% penicillin/streptomycin. After 7 days, we performed a K^+^ influx assay. We kept human SGCs proliferating beyond passage 12 from a DRG donor (NDRI). These SGCs were transfected with WT or *mutant GPR37L1* using TransIT®-LT1 reagent (Mirus Madison, WI, USA, Cat Bo.MR2300). The control and mutant GPR37L1 or KCNJ3 expression levels were examined by Real-time PCR.

### Preparation of plasma membrane (PM) fraction

Plasma membrane (PM) fraction was prepared from mouse or human DRG tissues and HEK cells using Mem-PER Plus Kit (Thermo Scientific, Cat No. 89842) and manufacture recommend protocol after biotinylation/streptavidin-magnetic bead (Thermo Scientific, Cat No.88817) pull down. Tissues or cells were harvested and washed out using a cold cell washing buffer. The tissues/cells were lysed by cell permeabilized buffer containing protease inhibitor (Sigma, Cat No. P8380) and phosphatase inhibitor (Sigma, Cat No.524631) using mechanical dissociation and ultrasonication. The lysate was centrifugated and the pellet was resuspended with PER lysis buffer containing protease and phosphatase inhibitor cocktail (Sigma).

### Western blot

Protein samples were prepared from transfected cells, lipid pull-down beads, and DRGs. Tissue and cells were placed on ice and lysed with ice-cold RIPA buffer (Sigma, Cat No. R0278) with Protease Inhibitor Cocktail Tablet (pH 7.4) (Roche Diagnostics, Cat No. 5892988001). The cell lysates were centrifugated to remove insoluble debris and the protein level was detected via BCA assay. The supernatant was mixed with 4x Laemmli buffer (BioRad, Cat No. 1610747) and boiled for 10 min (100 ℃, containing BME for reducing Bis-Tris PAGE). Protein samples were electroporated on 4-20% gradient SurePAGE™ Bis-Tris gel (Genscript) or 4-20% Tris-glycine gel (BioRad) and blotted on a PVDF membrane (BioRad). Ponceau S staining was used for the detection of total proteins. The primary antibody was incubated with 1% BSA at 4℃ overnight. We used the following primary antibodies: anti-GPR37L1 antibody (Bioss, Rabbit, 1:1000, Cat No. bs-15390R), anti-Kir4.1 antibody (Thermo Scientific, rabbit, 1:1000, Cat No. 12503-1-AP), Kir3.1/KCNJ3 antibody (Santacruz, mouse, 1:500, Cat No. sc-365457), anti-TfR antibody (Santa Cruz, mouse, 1:500, Cat No.sc-32272), anti-flag antibody (Cell signaling, Rabbit, 1:1000, Cat No.14793), and anti-GAPDH antibody (Proteintech, mouse, 1:2000, Cat No. 60004). Blots were further incubated with an HRP-conjugated secondary antibody (Santacruz, mouse, 1:2000, Cat No. sc-2357) or m-IgG Fc BP-HRP (Santacruz,1:5000, Cat No sc-525409) developed in ECL solution (Pierce). Some blots were also incubated with an Alexa Fluor® 647-conjugated donkey anti-rabbit IgG (Jackson ImmunoResearch,1:1000, Cat No. 711-605-152) or with Alexa Fluor® 647-conjugated donkey anti-guinea pig (Jackson ImmunoResearch,1:1000, Cat No. 706-605-148) and visualized in ChemiDoc XRS (BioRad) using chemiluminescence or Cy5 detection protocol. Protein signal intensity was quantified by Image Lab 6.1 (BioRad).

### Flow cytometry

Mouse DRG tissues were dissociated with 1 mg/ml Collagenase/Dispase (Roche) in a shaking incubator for 90 min. The dissociated tissues were incubated in 10% FBS-supplemented DMEM media for 1 h for neutralizing of the enzymes. The dissociated cells were washed out using a PBS + 10 mM EDTA solution. The cells were then fixed with 2% paraformaldehyde (PFA) and permeabilized with HBSS + 2% triton X-100. All dissociated cells were blocked with Fc receptors staining buffer (1% anti-mouse CD16/CD32, 2.4 G2, 2% FBS, 5% NRS, and 2% NMS in HBSS; BD Bioscience) and then stained with a standard panel of antibodies: Glast-PE (rat-IgG, Cat No.130-118-483, 1:200, Miltenyl Biotech), Nissle-cy5 (Sigma, 1 µg/ml), and GPR37L1-Apc-cy7 (rabbit IgG, Cat No. bs-15390R, Bioss, 1 µg/ml). After staining, cells were washed in PBS with EDTA. The flow cytometry events were acquired in a BD FACS Canto II flow cytometer by using BD FACS Diva 8 software (BD Bioscience). Data were analyzed using Cytobank Software (https://www.cytobank.org/cytobank).

### Computer simulations

The protein sequence of human GPR37L1 was downloaded from the UniProt database (ID: O60883) in *fasta* format. The predicted topology for GPR37L1 in UniProt was used for seven-transmembrane alignments and long-loop identification. Template selection and homology modeling were performed using the automated modeling server GPCR-ModSim (66). Human EDNRB (PDB: 6IGK) was selected as a template and active conformation of the model was generated. All other loops were also refined by the Prime module. Ligands were drawn in the Maestro suite in 2D format and were structurally preprocessed using LigPrep from the Schrodinger Suite (Schrödinger Release 2018-4: LigPrep, Schrödinger, LLC, New York, NY, 2018.). Protonation at a physiological pH (7±2) and energy minimization were performed using the OPLS3 force field. To elucidate the binding mode of all ligands in the binding site of the homology model of hGPR37L1, docking studies were performed with the help of Autodock4 software (67). Before the docking, the hGPR37L1 structure was prepared using the AutoDock Tools 4 software. The stability and intra-molecular conformational changes of the protein and molecular dynamics simulations (MDS) were performed on a 1000 ns time scale for the protein-ligand complex and 1000 trajectory structures were recorded. Using the GPU-accelerated DESMOND software (68), the top-scored docking poses were subjected to solvent-explicit, all-atom MDS. The OPLS-2005 force field was used for model generation of the protein-ligand complex, energy minimization, and MDS. The protein was inserted into the POPC lipid bilayer and the full system was immersed in a periodic orthorhombic water box TIP3P. The NPT ensemble class was used with the temperature set to 300 K and pressure set to 1.01325 bar. The trajectory clustering method of Desmond was used to cluster 1000 trajectory structures into ten clusters based on atomic root mean square deviation (RMSD). Then 100 ns MDS was performed on the cluster 1 structure to assess the stability of the docking complex. 2D structure was plotted using protter software.

### Prediction of protein stability

To predict the stability of the protein in mutant GPR37L1, we used web-based protein stability software in known or newly discovered mutations. Change instability from a mutant (M) protein to a wild-type (W) form was defined as the difference in the corresponding unwinding free energies ΔΔG of the two proteins. A negative ΔΔG value indicates that the mutant protein is more unstable. To calculate the protein stability, we used 9 different web-based prediction servers and presented the ΔΔG value(69, 70). The prediction servers are as follows:

PremPS: https://lilab.jysw.suda.edu.cn/research/PremPS/

DDgun: https://folding.biofold.org/ddgun/index.html#!

Mupro: http://mupro.proteomics.ics.uci.edu/

SAAFEC-SEQ: http://compbio.clemson.edu/SAAFEC-SEQ/

I-mutant: https://folding.biofold.org/cgi-bin/i-mutant2.0.cgi

DUET: http://biosig.unimelb.edu.au/duet/stability

DeepDDG: http://protein.org.cn/ddg.html

mCSM: http://structure.bioc.cam.ac.uk/mcsm

SDM: http://www-cryst.bioc.cam.ac.uk/~sdm/sdm.php

### In situ hybridization using RNAscope probes

Mice were transcranial perfused with PBS followed by 4% PFA under deep anesthesia with isoflurane. Lumbar DRGs, trigeminal ganglia, and nodose ganglia were isolated and post-fixed in the same fixative. Tissues were cryopreserved in a sucrose gradient, then embedded in OCT medium (Tissue-Tek), and cryosectioned at 14 μm. The RNAscope probes against mouse *Gpr37l1* (Cat No. 319301), mouse *Gpr37* (Cat No. 319291), human *GPR37L1* (Cat No. 513641), human *KCNJ3* (Cat No.1154801-C3), human *KCNJ10* (Cat No.461091), and human *TUBB3* (Cat No.481251) were designed by Advanced Cell Diagnostics and the RNAscope multiplex fluorescent assays we conducted according to the manufacturer’s instructions. Pre-hybridization and hybridization were performed according to standard methods (34).

### Immunohistochemistry

Mice were deeply anesthetized with isoflurane and perfused through the ascending aorta with PBS, followed by 4% PFA. After the perfusion, L4-L5 DRGs were removed from the mice and post-fixed in the same fixative overnight. Human DRGs were obtained from the National Disease Research Interphase (NDRI) and fixed in 4% PFA overnight. The samples were then dehydrated with a 10% to 30% sucrose gradient, embedded in Tissue-Tek O.C.T., and sliced into sections (14 μm) in a cryostat. The sections were blocked with 5% donkey serum and 0.2% triton X-100 for 1 h at room temperature, then incubated overnight at 4 °C with the following primary antibodies: anti-GPR37L1 antibody (Alomone Labs, Rabbit, 1:500, Cat No. AGR-050), anti-GS antibody (Novus Biologicals, Rabbit, 1:1000, Cat No. NBP2-32241), anti-FABP7 (Neuromics, Mouse,1:1000, Cat No.MO22188), and anti-KCNJ10/Kir4.1 antibody (Thermo Scientific, Guinea pig, 1:800, Cat No. PA5-111798). The sections were washed in PBS and incubated with the following secondary antibodies (1:500, Biotium) for 2 h at room temperature: CF633-donkey anti-rabbit (Cat No. 20125), CF568 donkey anti-guinea pig (Cat No. 20377), and CF488A donkey anti-mouse (Cat No. 20952). For clarity, channel colors were exchanged in the presented mouse DRG images without disrupting the signal using ImageJ. DAPI (1:1,000, Thermo Scientific, Cat No. 62248) was used to stain the cell nuclei in tissue sections. The sections were then washed with PBS, mounted in Fluoromount G mounting medium (Southern Biotech, Cat No. 0100-01), and observed under a confocal laser scanning microscope (SP5 Inverted confocal-LSRC, Leica Microsystems). Some stained sections were also examined with a Leica SP5 or Zeiss 880 confocal microscope with Z-stack and Tile Scan. The maximum projected and stitched images were produced using the Zeiss Zen software or with ImageJ.

### Patch-clamp recordings in mouse SGCs in whole-mount DRG preparation

Under urethane anesthesia, mice were rapidly euthanized, followed by careful isolation of lumbar DRGs placed in the oxygenated artificial cerebral spinal fluid (ACSF). DRGs were briefly digested (20 min) using an enzymatic mixture consisting of collagenase A (1 mg/mL) and Trypsin (0.25% original solution). Intact DRGs were then incubated in ACSF oxygenated with 95% O_2_ and 5% CO_2_ at 34°C. Following incubation, DRGs were transferred to a recording chamber and continuously perfused (∼3 ml/min) with ACSF. SGCs in whole mouse DRG could be visualized using a 40x water-immersion objective on an Olympus BX51WI microscope. The round or fusiform-shaped cell bodies of SGCs have small sizes (<10 μm) but are visible near the edges of DRG neurons. Patch pipettes (Chase Scientific Glass Inc.) were pulled and filled with a pipette solution containing (in mM): 126 potassium gluconate, 10 NaCl, 1 MgCl_2_, 10 EGTA, 2 Na-ATP, and 0.1 Mg-GTP, adjusted to pH 7.3 with KOH. The resistance of pipettes was 10-12 MΩ. A Whole-cell patch-clamp configuration was made on SGCs at room temperature using a Multiclamp 700B amplifier (Axon Instruments, Union City, CA). Under voltage clamp, at a holding potential of -80 mV, inward or outward currents were triggered by voltage steps beginning at -160 mV and increasing by 20 nM every 200 ms to a maximum of +40 mV (71). To isolate the inwardly rectifying potassium current (Kir), potassium currents from the same cells were recorded in the absence and presence of 100 μm extracellular barium for the blockade of the Kir4.1 channel (35, 72). The Kir4.1 currents were obtained by digitally subtracting those currents in the absence and presence of barium.

### Ti^+^ influx assay

Ti^+^ flux assay was conducted using the manufacture’s and standard protocol (73). For examining Ti^+^ flux in GPR37L1, KCNJ3, KCNJ3/KCNJ10-expressing Hek293 cells, and WT/E296K-GPR37L1-expressing human SGCs, cells were dislodged from tissue culture flasks using TrypLE Express reagent (Gibco, Cat No. 12604013) and transferred to a 50 mL centrifuge tube, and centrifuged at 500g for 2 min. The supernatant solution was removed by aspiration, and the pellet was resuspended at a concentration of ~ 1000 cells/μL in a cell culture medium. Then 100 μl/well of the cell suspension was transferred to 96-well plates. The seeded cells were incubated overnight in a humidified 5% CO2 cell culture incubator at 37 °C in low FBS media (2%). After overnight incubation, the cell culture medium was replaced with thallium influx dye-loading solution consisting of assay buffer (Hanks Balanced Salt Solution + 10 mM HEPES), 0.04% (w/v) Pluronic F-127 (Sigma), 1 μM of Thallos-AM (Ion bioscience) and 1 x TRS solution. Following a 60 min incubation at room temperature, the dye-loading solution was replaced with HBSS + 2 mM Ca^2+^ and + 2 mM Mg^2+^ 90 µl/well of assay buffer, and the plates were loaded into an Infiniti-M200 pro reader (Tecan, Männedorf, Switzerland). Data were acquired at (excitation 480, emission 520 nm) for 30 s intervals, followed by the addition of 10 μL/well of test compounds (30 min for KCNJ3 cell line or 1 h for KCNJ10 cell line). After baseline measurement for 5 min, 10 μl/well Ti^+^ stimulation was injected (at a rate of 100 μl/sec), and fluorescence was measured for 10 min. The Tl^+^ stimulus buffer consisted of 125 mM KHCO_3_, 1.8 mM CaSO_4_, 1 mM MgSO_4_, 5 mM glucose, 5 mM Tl_2_SO_4_, 10 mM HEPES, pH 7.4. The cover-glassed cultures of mouse and human SGCs were imaged using an Orca6 camera (Hamamatsu) every 3 sec with excitation at 480 nm and emission at 520 nm filters after thallium influx dye-loading solution. The imaging was performed using the MetaFluor software (Molecular Devices) in HBSS buffer supplemented with 2 mM Mg^2+^ and 2 mM Ca^2+^. The activity was measured following stimulation with 500 µM Ti^+^ solution.

### Nociceptive behavior tests

*Gpr37l1*^+/+^, *Gpr37l1*^+/-^, or *Gpr37l1*^-/-^ mice were habituated to the testing environment for at least two days before baseline testing. All animal behaviors were tested blindly. Thermal and mechanical sensitivity was tested before and after the injection of paclitaxel (PTX, 1 x 6 mg/kg, or 4 x 2 mg/kg, IP) or streptozotocin (STZ, 75 mg/kg, IP). MaR1 was intrathecally injected 3 days after the STZ or PTX injection. For testing mechanical sensitivity, mice were confined in boxes (14 × 18 × 12 cm) placed on an elevated metal mesh floor, and their hind paws were stimulated with a series of von Frey hairs with logarithmically increasing stiffness (0.16-2.00 g, Stoelting), presented perpendicularly to the central plantar surface. The 50% paw withdrawal threshold was measured by Dixon’s up-down method (74). Thermal sensitivity was measured using a Hargreaves radiant heat apparatus (75) (IITC Life Science). The basal paw withdrawal latency was adjusted to 10-15 s, with a cutoff of 25 s to prevent tissue damage. Acetone (50 μl) was applied through wire mesh flooring onto the plantar surface of the infected hindpaw to produce evaporative cooling (59). Mechanical and thermal sensitivity was also tested after intra-DRG injection of siRNA and MaR1.

### ELISA assay

Mouse IL-1β levels were measured using a mouse IL-1β ELISA Kit (R&D, Cat No. MLB00C, Minneapolis, MN), and human IL-1β levels were detected using a human IL-1β ELISA Kit (Proteintech, Cat No. KE00021) according to the manufacturer’s instructions. Briefly, 100 μl of culture medium were added to each well and a standard curve was included for each experiment.

### Real-time quantitative PCR

Real-time quantitative PCR (RT-qPCR) assays were conducted in cDNA samples obtained from a reverse-transcription reaction, using the BioRad CFX96 system (BioRad). Total RNA from the DRG and SGC cultures was extracted using the Direct-zol RNA Miniprep Kit (Zymo Research), and 0.5–1 μg RNAs were reverse transcribed using the iScript cDNA Synthesis Kit (Bio-Rad). Specific primers, including the GAPDH control, were designed using IDT SciTools Real-Time PCR software. We performed gene-specific mRNA analyses using the CTX96 Real-Time PCR System (Bio-Rad). Quantitative PCR amplification reactions contained the same amount of reverse transcription product (50 ng), including 7.5 μl of 2× iQSYBR Green Mix (Bio-Rad) and 100–300 nM forward and reverse primers, in a final volume of 15 μl. The primer sequences are listed below. Primer efficiency was obtained from the standard curve and integrated for the calculation of relative gene expression, which was based on real-time PCR threshold values of different transcripts.

*mGpr37l1* forward: *GCATTGTGTGGCACAGCTAC*

*mGpr37l1* reverse: *AAACTGGAAATGCCAGCGTG*

*mKcnj10* forward: *TGCGGAAGAGTCTCCTCATTGG*

*mKcnj10* reverse: *GTCTGAGGCTGTGTCTACTTGG*

*mGapdh* forward: *AGGTCGGTGTGAACGGATTTG*

*mGapdh* reverse: *GGGGTCGTTGATGGCAACA*

*hGPR37L1* forward: *ACGAGATCACCAAGCAGAGGCT*

*hGPR37L1* reverse: *GCACAGAGGCTGAAAGTCGTGA*

*hKCNJ3* forward: *GATCTCCATGAGGGACGGAAAAC*

*hKCNJ3* reverse: *GAAGGAACTCACCCTCAGGTGT*

*hGAPDH* forward: *GTCTCCTCTGACTTCAACAGCG*

*hGAPDH* reverse: *ACCACCCTGTTGCTGTAGCCAA*

### Genome-wide association

Genome-wide association tests were performed in the large UK Biobank project cohort comprised of half a million participants (41, 76). Information about chronic pain reports collected at the visit was available at eight body sites: head (headaches), face, neck/shoulder, stomach/abdominal, back, hip, knee, and widespread. Control subjects were those that answered, “none of the above” at field 6159 for the question “In the last month have you experienced any of the following that interfered with your usual activities?”. Case subjects for chronic pain at body site X (among the eight listed) were those that answered “yes” to the question “Have you had pains for more than 3 months?”; fields are 3571 (back), 4067 (face), 2956 (widespread), 3799 (headaches), 3414 (hip), 3773 (knee), 3404 (neck/shoulder), and 3741 (stomach/abdominal). Input genotyping data for association tests were from whole-exome sequencing (in VCF format), available for 200K individuals. Variants were annotated to GRCh38 reference provided by UK Biobank (http://biobank.ndph.ox.ac.uk/ukb/refer.cgi?id=3803) with Ensembl Variant Effect Predictor (VEP, release 99)(77). Rare coding variants were tested, and we required a minor allelic frequency of less than 0.01 and a minor allele count greater than 5. We used SAIGE (version 0.44.2) (78) to perform the tests, as it considers cryptic relatedness and guards against false-positive associations in the face of case-to-control imbalance (due to the cross-sectional nature of the cohort) by means of the saddle point approximation method (79). Covariables were: age, age squared, sex, genotyping platform, recruitment centers, and the first 40 principal genetic components. Subjects were of White British origin (field 22006). Discarded subjects were those that opted out of the study, failed genotyping or imputation quality controls, or displayed sex aneuploidy or mismatched declared / genetically-determined sex. Retained SNPs displayed at least five counts of the minor allele in selected cases or control subjects. The meta-analysis at a given variant position across all eight chronic pain sites was performed using the inverse variance-based weighting scheme (80). A combined Annotation Dependent Depletion score (CADD v1.6) (43) was used to estimate the deleteriousness of the variants. It computed PHRED-scale CADD scores for variants, which provided a relative order compared to all existing variants in the human genome. Higher CADD scores for higher risk of deleteriousness.

### Statistics

All data were expressed as the mean ± SEM. The sample size for each experiment was indicated in figure legends. GraphPad Prism 8.0 software was used to perform statistical analysis. The data were analyzed by two-way ANOVA, followed by Bonferroni’s post-hoc test or Tukey’s post-hoc test for multi-group comparison, or by Student t-test for two-group comparison. *P*<0.05 was considered statistically significant. Statistical significance was indicated as * *P*<0.05, ** *P* <0.01, *** *P* <0.001, **** *P* <0.0001.

### Study Approval

All the animal procedures were approved by the Institutional Animal Care & Use Committee of Duke University. Animal experiments were conducted in accordance with the National Institutes of Health Guide for the Care and Use of Laboratory Animals.

## Data and Software Availability

No custom software was used in this study. The data are available upon reasonable request.

## Supplemental Information

Supplemental information includes 10 supplemental figures and 5 supplemental tables.

## ACKNOWLEDGMENTS

This study was supported by Duke University Anesthesiology Research Funds, NIH R01 grants DE17794 and 1NS13181201A1, and DoD grants W81XWH2110885 and W81XWH2110756. L.D. was supported by a professorship in Pain Research from Pfizer Canada, the Canadian Excellence Research Chairs grant CERC09, and NIH grant U54 DA049110. T.B. was supported by NIH R01 grant NS113243. Part of this study was conducted under UK Biobank application no. 20802.

## AUTHOR CONTRIBUTIONS

S.B. and R.R.J. developed the project. S.B, C.J., J.X., S.C., A.M., X. L., Q.H, Y.L. and Z.W. conducted experiments and data analyses; X.O., M. P., L. O. F, and S. J. E. contributed to genome association study under the supervision of L.D. R.T., T.B., and Q.Z. participated in project development. S.B. and R.R.J. wrote the manuscript; L.D., T. B., and other co-authors edited the manuscript.

---

**Supplemental Table 1:** Total GPCR transcripts (A) and orphan GPCR transcripts (B) in mouse DRGs.

**Supplemental Table 2:** Total GPCR transcripts (A) and orphan GPCR transcripts (B) in human DRGs.

**Supplemental Table 3:** (A) SAIGE association tests for rare coding variants and a report of chronic pain at eight body sites. Columns are: “PainSite” for body site of chronic pain; “markerID” for variant ID; “Chr” for chromosome number of variant; “Pos” for GRCh38 genomic coordinate of variant; “Allele1” for the reference allele of variant; “Allele2” for the effective allele of variant; “rsid” for known rsid of variant; “AC2” for allele account of the effective allele; “AF2” for allele frequency of the effective allele; “AF2.Cases” and “AF2.Ctrls” for allele frequency of the effective allele in group cases and controls, respectively; “N.Cases”, “N.Ctrls”, and “N” for number of samples in group cases, controls, and total samples; “Consequence” for functional consequence predicted by VEP; “BETA” for effect size of effective allele; “SE” for standard error of “BETA”; “odds[95%]” for odds ratio and 95% confidence interval; “P” for raw P value; (B) Meta-analysis on SNP rs148475636 and rs767987863. Columns represent the same meaning as in Table 3A. Meta results are in row “Meta”.

**Supplemental Table 4:** Animals used in this study.

**Supplemental Table 5:** Human DRG donners used in this study. (A) Human DRGs from NDRI. (B) Information of human DRG donors with neuropathic pain (*n* = 3) and controls (*n* = 3).

**Supplemental Figure 1.**
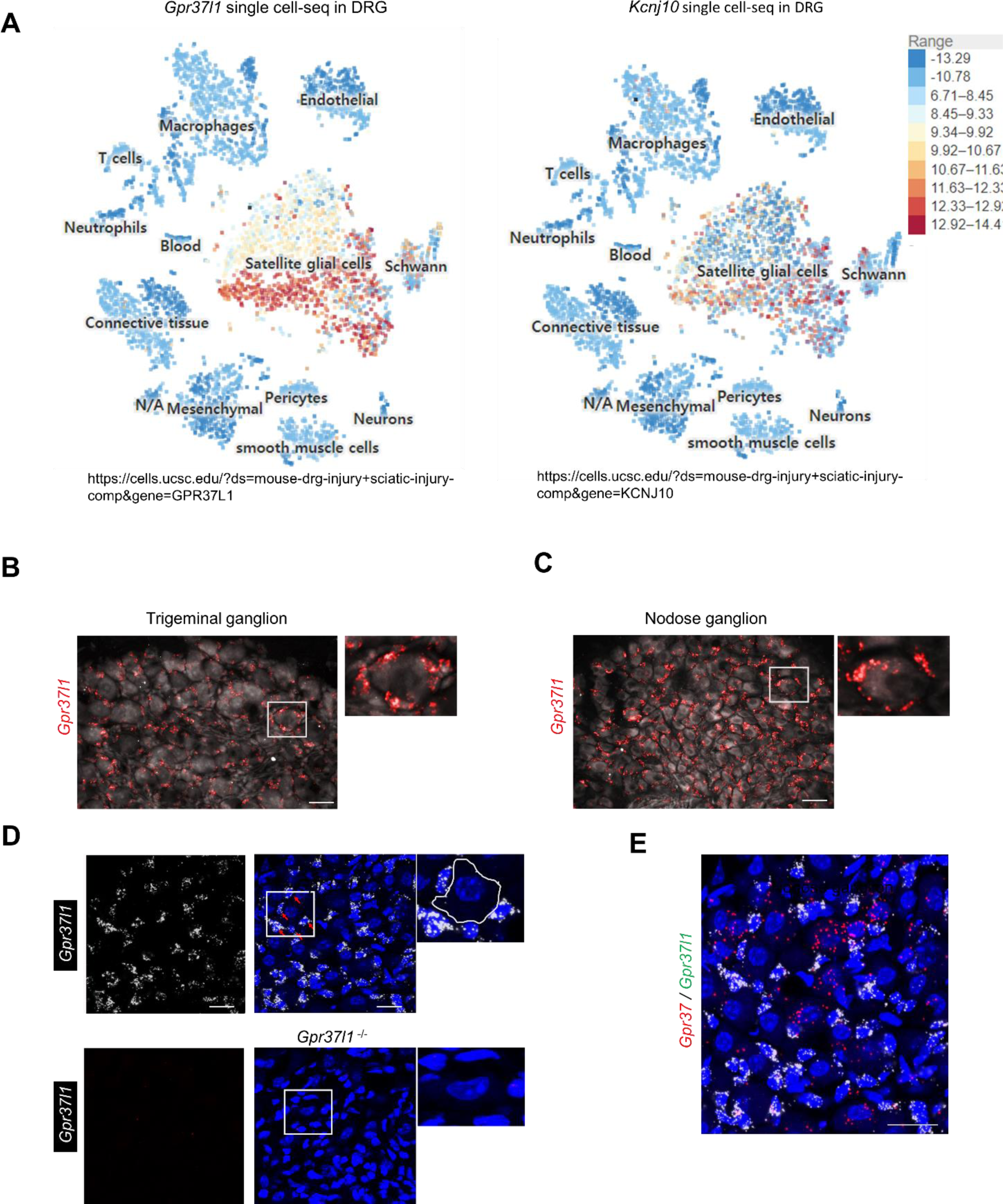
Mouse RNA sequencing and human microarray analysis. (**A**) Single-cell RNAseq showing expression of *Gpr37l1* and *Kcnj10* transcripts in SGCs of mouse DRG tissues (19). (**B, C**) Images of RNAscope in situ hybridization (ISH) of *Gpr37l1* expression in trigeminal ganglion (TG, B) and nodose ganglion (NG, C). Small boxes are enlarged in the right panels. Neurons are lightly labeled with Nissl staining (white). Scales, 25 µm. (**D**) RNAscope images of *Gpr37l1* expression in NG of *Gpr37l1* ^+/+^ mice (WT, top) and *Gpr37l1* ^-/-^ mice (bottom). Small boxes are enlarged in the right panels. Scales, 25 µm. (**E**) Double staining of RNAscope ISH showing distinct expression of *Gpr37* (red) and *Gpr37l1* (white) in mouse NG. Scale = 25 µm.

**Supplemental Figure 2.**
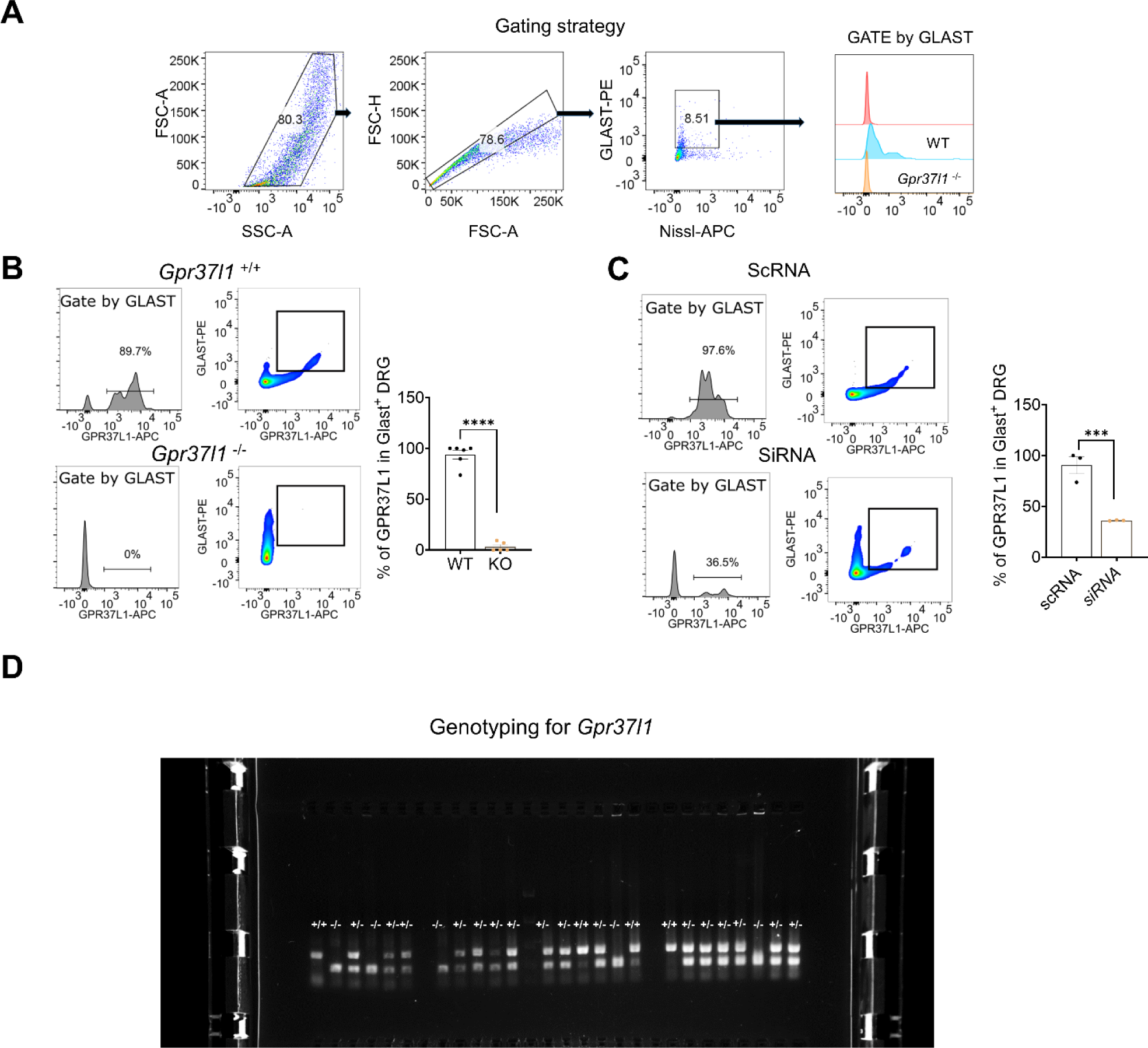
Characterization of GPR37 expression in mouse DRG tissue by flow cytometry. (**A-C**) Flow cytometry data showing GPR37L1 expression in DRGs of *WT* and *Gpr37l1* ^-/-^ mice. (**A**) Gating strategy. **(B)** GPR37L1 expression in DRGs of *WT* and *Gpr37l1* ^-/-^ mice. Left, representative images of flow cytometry and histograms after gating by Glast-PE. Right, Quantification of GPR37L1+ cells in the Glast-PE+ population (*n* = 6 for WT and *n* = 5 for KO mice). **(C)** Left, representative images of flow cytometry and histogram after gating by Glast-PE in DRGs of scRNA or siRNA treated mice (related to Figure 3G). Right, Quantification of GPR37L1+ cells in the Glast-PE+ population (*n* = 3 mice/group). GLAST was used as a marker to isolate SGCs in DRGs. (**D**) Genotyping for *Gpr37l1* WT and mutant mice. Data are expressed as mean ± s.e.m. \*\*\*\**P*<0.0001, \*\**P*<0.01, unpaired t-test.

**Supplemental Figure 3.**
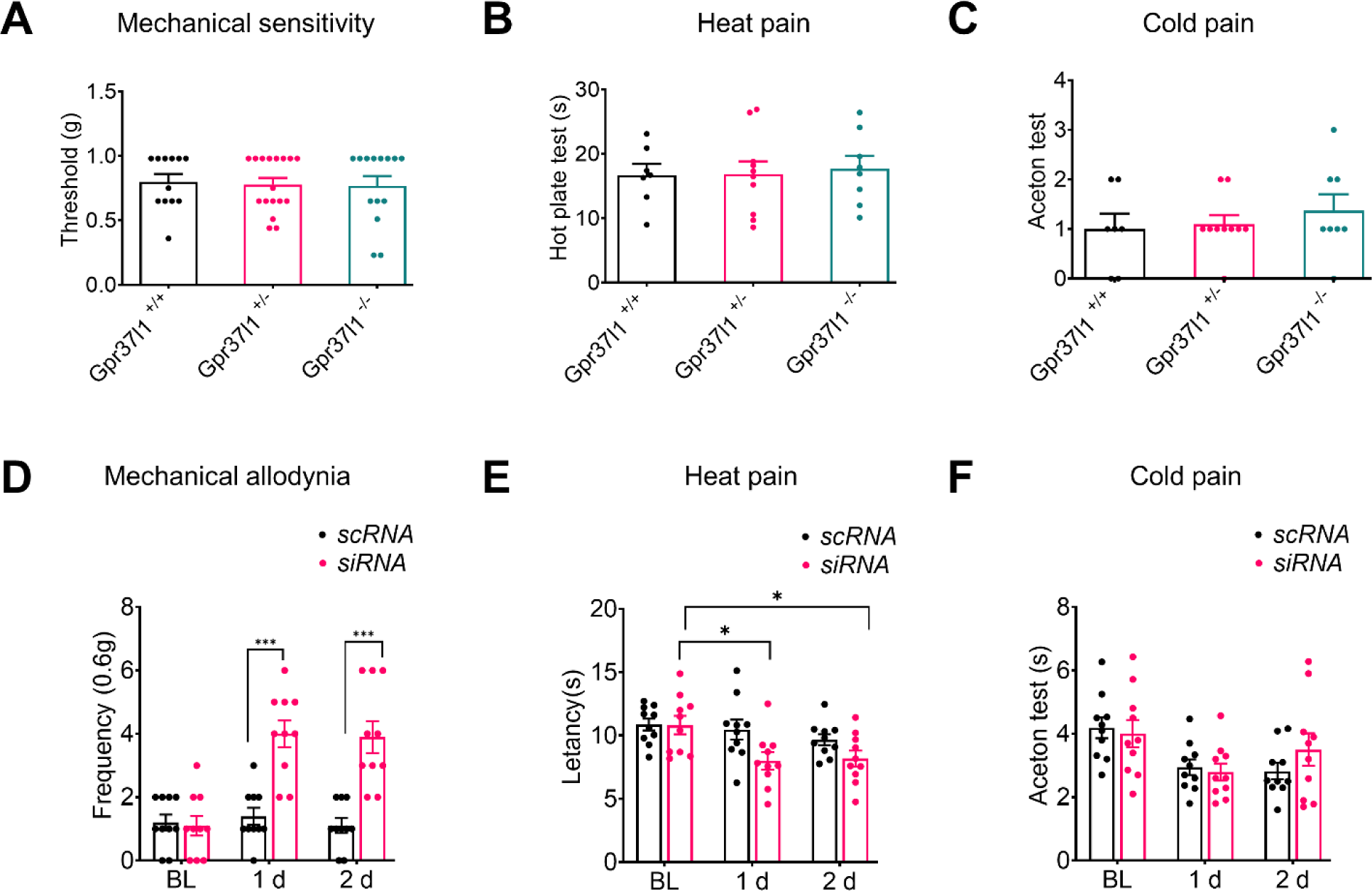
Characterization of mechanical and thermal pain in WT mice and *Gpr37l1* mutant mice. (**A-C**) Baseline pain sensitivity in *Gpr37l1* ^+/+^ mice (*n = 12*), *Gpr37l1* ^+/-^ mice (*n =* 12), *and Gpr37l1* ^-/-^ mice (*n =* 17), assessed in von Frey test (mechanical sensitivity, A), hot-plate test (heat sensitivity, B), and acetone test (cold sensitivity, C). (**D-F**) Mechanical and thermal hyperalgesia in mice treated with I.G. injection of *Gpr37l1* siRNA (*n =* 10) and control scRNA (*n =* 10). (**D**) Paw withdrawal frequency (0.6 g) showing the siRNA-induced mechanical allodynia at 1d and 2d. (**E**) Hargreaves test showing siRNA-induced heat hyperalgesia at 1d. (**F**) Acetone test shows no cold allodynia following the siRNA treatment. Data are expressed as mean ± SEM and statistically analyzed by One-Way ANOVA with Turkey’s post-hoc test (A, B), Kruskal-Wallis test with Dunn’s post-hoc test (C), or Two-Way ANOVA with Bonferroni’s post-hoc test (scRNA vs siRNA, D-F). \**P*<0.05, \*\*\**P*<0.001.

**Supplemental Figure 4.**
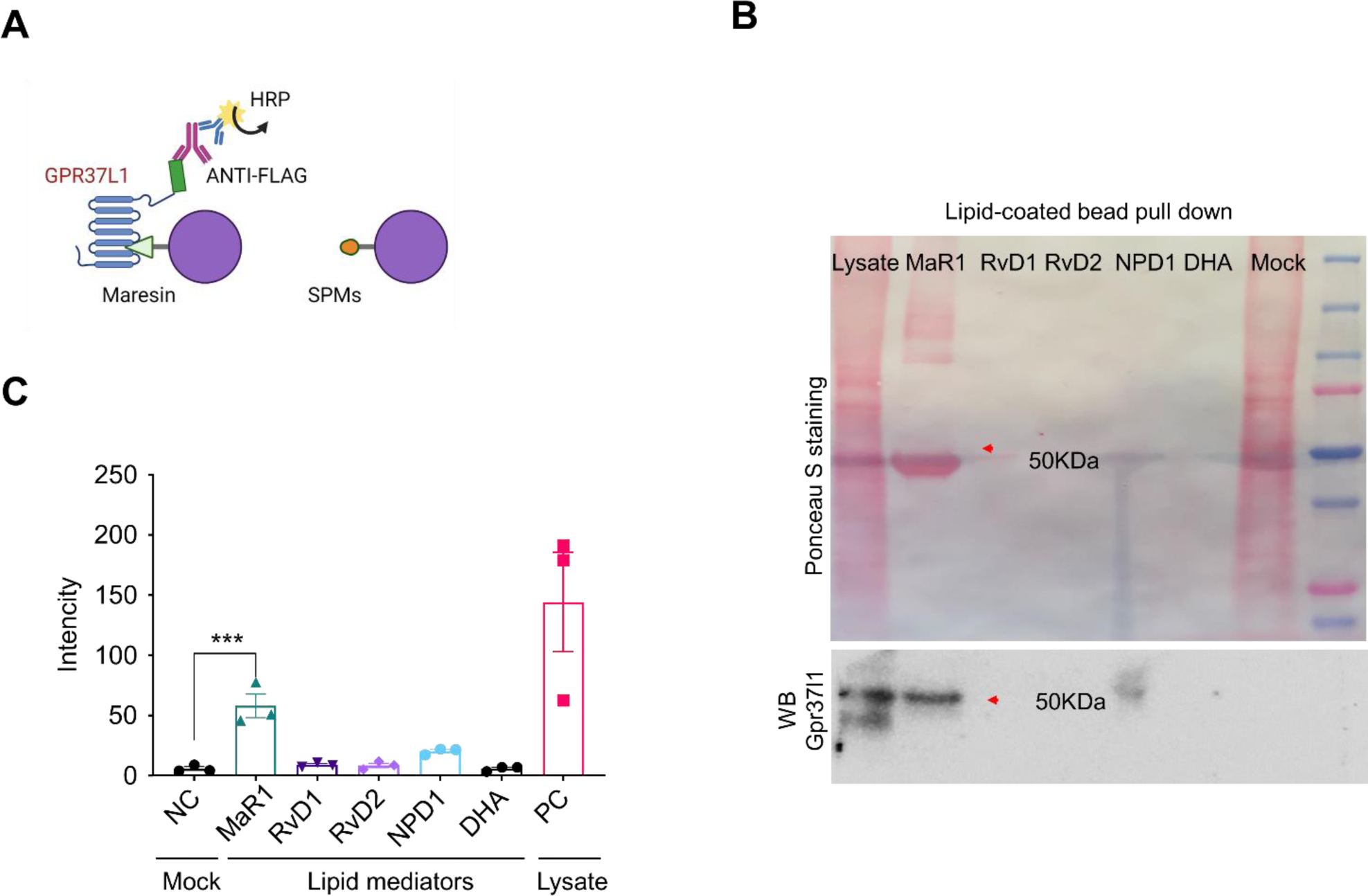
GPR37L1 binds with MaR1. Lipid pull-down assay showing MaR1 binding to GPR37L1 in GPR37L1-expressing HEK293 cells. **(A)** Schematic of the lipid pull-down assay using agarose bead-coated SPM (MaR1). **(B)** Top, Ponceau S staining; red arrowhead indicates the specific band at 50 kDa. Bottom: Anti-FLAG Western blot showing the GPR37L1 band, indicated by red arrowhead. One microgram of protein was loaded. (**C**) Quantification of the intensity of immune blots; DRG lysate and Mock were used as positive control (PC) and negative control (NC). *n* = 3 repeats. Data are expressed as mean ± s.e.m. and analyzed by One-Way ANOVA followed by Tukey’s post-hoc test (C). \*\*\**P*<0.001.

**Supplemental Figure 5.**
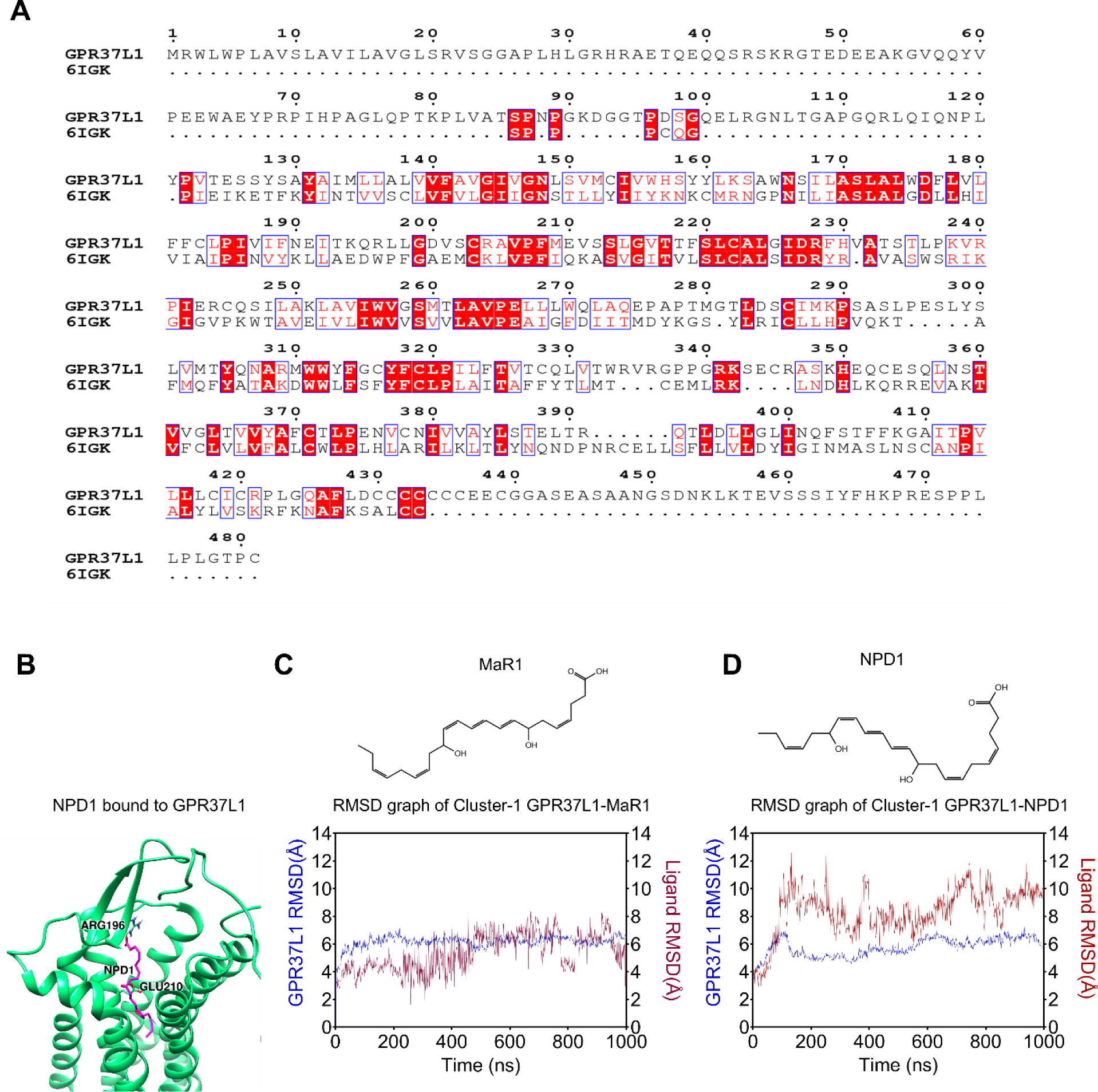
Computer simulation for the interaction of GPR37L1 and MaR1. (**A**) Alignment of amino-acid sequences of a/the crystallized construct of human EDNRB (PDB: 6IGK) and human GPR37L1 (UniProt ID: O60883). Conservation of the residues is indicated as follows: red panels for completely conserved; red letters for partly conserved; and black letters for not conserved. Note there is a 30.23% homology between these two genes. **(B)** The overall structure of hGPR37L1 (Green) complex with NPD1 (Magenta). **(C)** 1000 ns molecular dynamics simulation of the GPR37L1-MaR1 complex (red) or GPR37L1(blue). **(D)** 1000 ns molecular dynamics simulation of the GPR37L1-NPD1 complex (red) or GPR37L1(blue). Note that MaR1, but not NPD1, has stable interaction with GPR37L1.

**Supplemental Figure 6.**
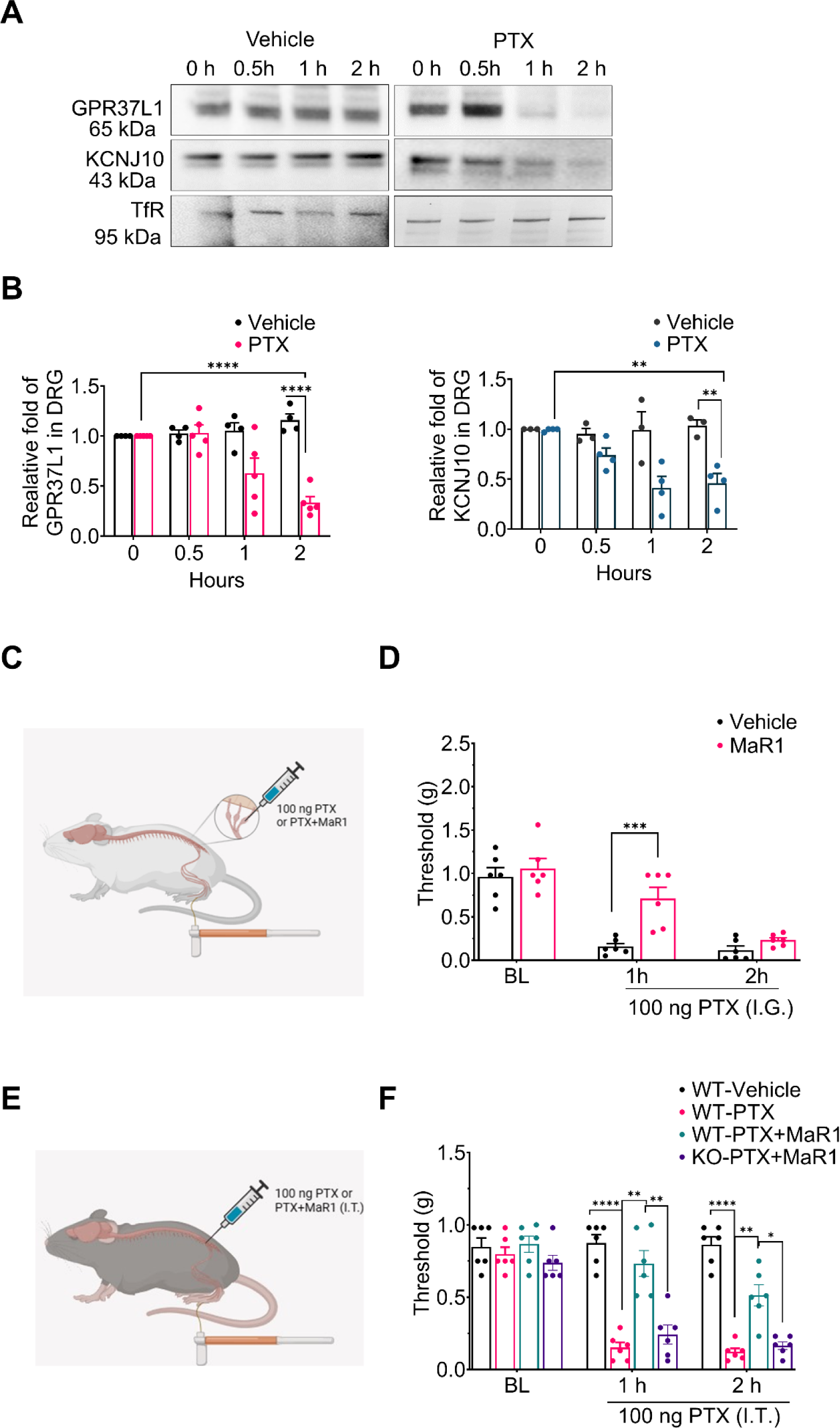
Chemotherapy causes rapid downregulations of plasma membrane (PM) expression of GPR37L1 and KCNJ10 in whole mount DRGs. **(A)** Time course of PM expression of GPR37L1 and KCNJ10 following treatment of vehicle (left) and paclitaxel (PTX, 1 μM, right). **(B)** Quantification of PM fraction of GRP37L1 (left) and KCNJ10 (right) expression. *n* = 3 for vehicle and *n* = 4 for PTX. **(C-F)** Unilateral local injection of paclitaxel (PTX, 100 ng) via I.G. route (C) or I.T. route (E) is sufficient to induce acute mechanical pain. **(C-D)** MaR1 (I.G., 100 ng) prevented PTX (I.G.) induced acute mechanical allodynia. **(E-F)** MaR1 (I.T., 100 ng) attenuated PTX (I.T.) induced acute mechanical pain in WT mice but not *Gpr37l1 ^-/-^* (KO) mice. *n* = 6 mice/group. Data are expressed as mean ± s.e.m. and analyzed by Two-Way ANOVA followed by Tukey’s post-hoc test (B, F) and Bonferroni’s post-hoc test (D). \**P*<0.05, \*\**P*<0.01, \*\*\**P*<0.001, \*\*\*\**P*<0.0001.

**Supplemental Figure 7.**
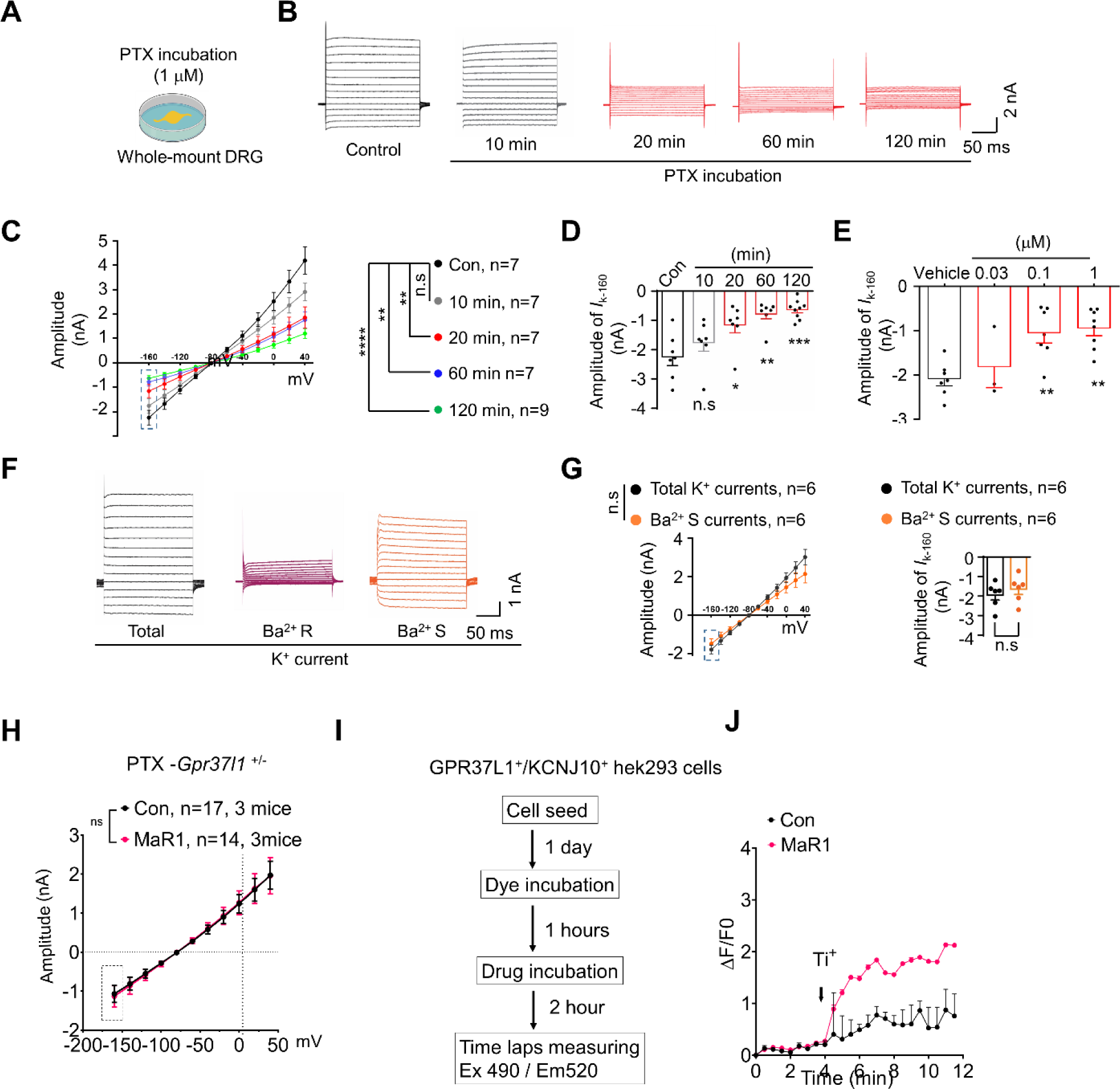
Characterization of K^+^ currents in SGCs of whole mount DRGs and Thallium (Ti^+^) influx assay in HEK293 cells. **(A)** Schematic diagram for whole-mount SGC recording. **(B)** K^+^ current traces in SGCs at different times after PTX treatment (1 μM, 10 min 20 min, 60 min, and 120 min) were recorded using the step voltage injection protocol. **(C)** Voltage-current curves in SGCs at different times after PTX treatment. **(D)** Quantification of the amplitude of -160 mV currents in (B) and (C). **(E)** Quantification of the amplitude of -160 mV currents at different doses of PTX (vehicle, 0.03 *n* = 3, 0.1,1 µM, *n* = 7 cells) at 20 min. **(F-G)** Isolation of Barium sensitive currents in SGCs in whole-mount DRGs. **(F)** Example traces of Barium sensitive currents in SGCs. **(G)** Left, average voltage-current curves in SGCs for Barium sensitive or insensitive currents (*n =* 6 cells, left). Right, quantification of -160 mV currents for Barium sensitive and total currents (*n =* 6 cells, right). (**H**) Average I/V curves in *Gpr37l1*^+/-^ SGCs after treatment of PTX *(n =* 17 cells) or PTX + MaR1 (*n* = 14 cells). **(I-J)** 96well plate Ti^+^ influx assay in GPR37L1 and KCNJ10 expressed HEK293 cells after MaR1 incubation. **(I)** Flowchart of Ti^+^ assay. **(J)** Representative traces of Ti^+^ influx in cells treated with 100 nM MaR1 (red, *n* = 3) or control vehicle (Con, black, *n* = 3) for 1h incubation. Data are expressed as mean ± SEM and analyzed by Two-way ANOVA with Tukey’s post-hoc test (C, G left), One-way ANOVA with Tukey’s post-hoc (D, E), and unpaired t-test (H). \**P*<0.05, \*\**P*<0.01, \*\*\**P*<0.001, **** *P*< 0.0001, n.s., not significant.

**Supplemental Figure 8.**
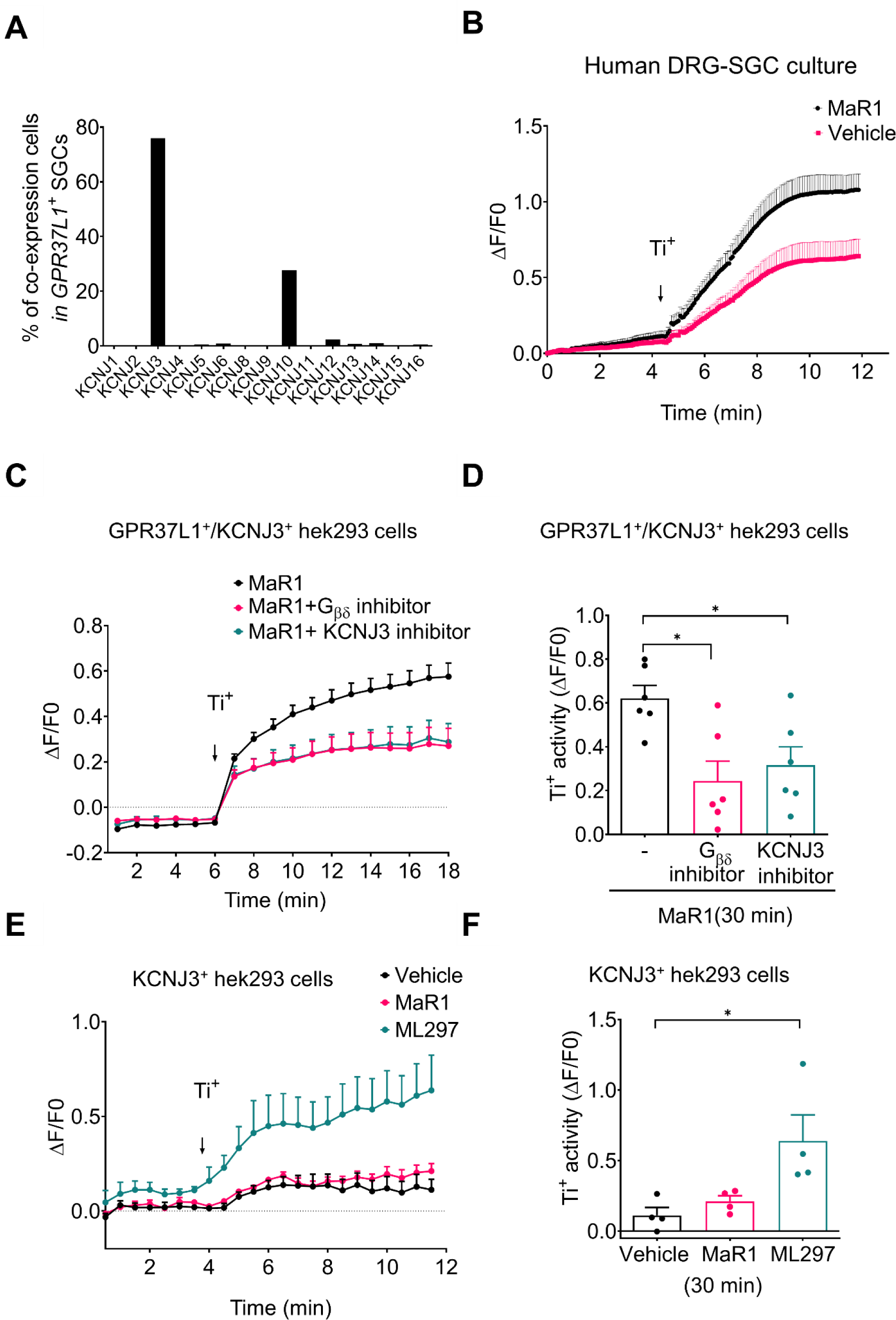
GPR37L1 regulates KCNJ3-mediated K^+^ influx in human DRG and HEK293 cells. **(A)** Co-expression percentage of *KCNJ3* and *KCNJ10* in GPR37L1+ SGCs from a human TG database of RNAseq (36). **(B)** Representative traces for Ti^+^ influx assay of cultured human SGCs treated with MaR1 (100 nM, *n* = 7) or vehicle (*n* = 8). **(C-D)** Ti^+^ influx assay in GPR37L1 and KCNJ3 expressing HEK293 cells. MaR1 increased Ti^+^ influx, which was reduced by Gβγ inhibitor (Gallein, 1 µM) and KCNJ3 inhibitor (SCH 23390, 1 µM). The baseline value of the Ti^+^ influx of the vehicle group was subtracted. **(C)** Traces of time-dependent Ti^+^ influx and the effects of Gβγ or KCNJ3 inhibitors (*n* = 14 at 30 min of drug incubation). **(D)** Quantification of Ti^+^ influx at 10 min *(n* = 6 culture). (**E-F**) Thallium influx assay in KCNJ3-expressing HEK293 cells. **(E)** Time course of Ti^+^ influx activity after vehicle, MaR1 (30 nM), and ML297 (300 nM). *n* = 4 cultures. **(F)** Quantification of Ti^+^ influx at 10 min *(n* = 4 cultures). Data are expressed as mean ± SEM and analyzed by One-way ANOVA with Tukey’s post-hoc (D, F) \**P*<0.05.

**Supplemental Figure 9.**
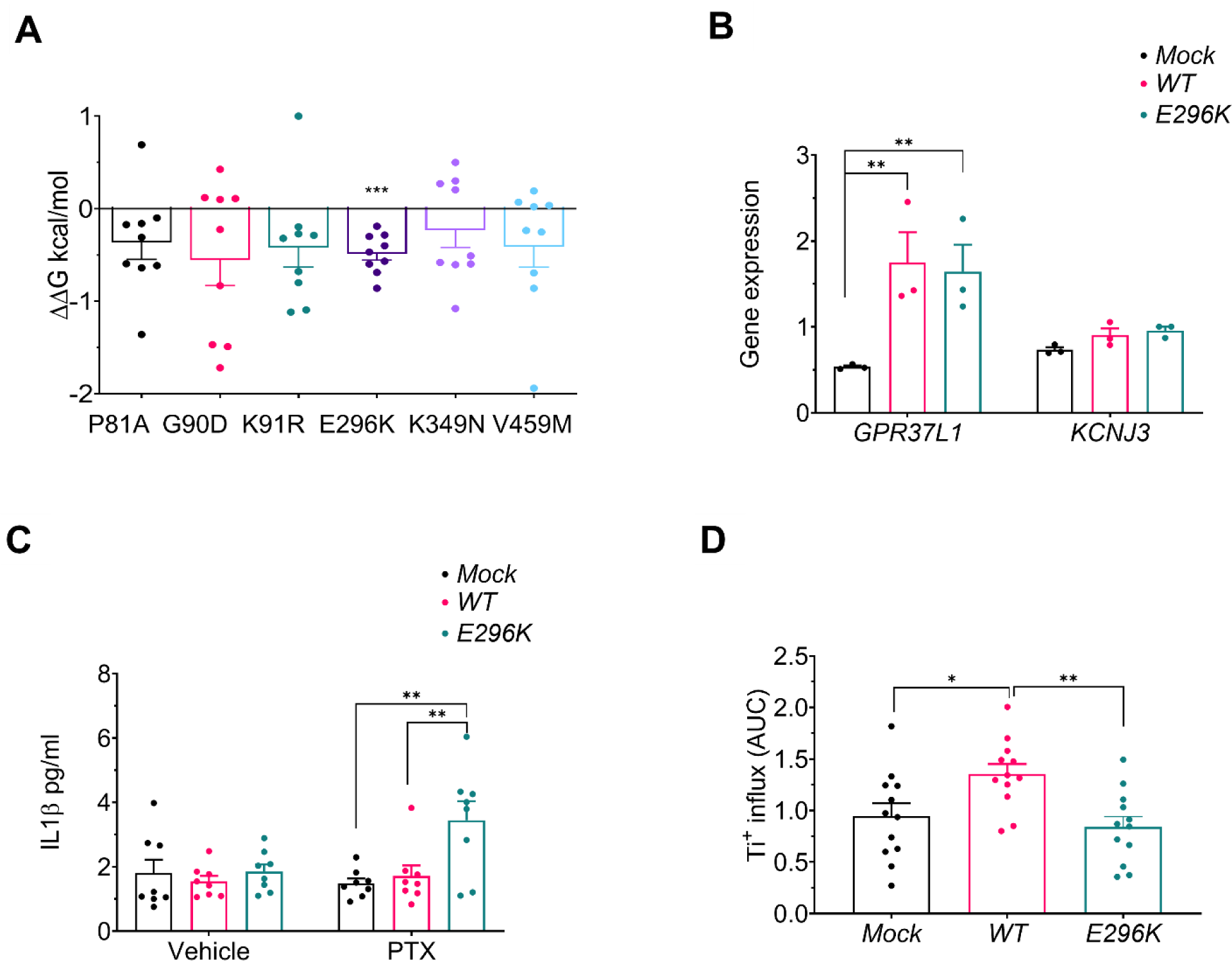
Prediction of GPR37L1 mutant stability and effects of E296K mutation on GPR37L1 and KCNJ3 expression, IL-1β release, and Ti^+^ influx. **(A)** GPR37L1 stability changes in different *GPR37L1* mutations were tested by protein stability prediction server using 9 different algorithms. **(B)** Realtime PCR shows the expression levels of *GPR37L1* and *KCNJ3* in mock, WT, or *E296K* mutant transfected human SGC cultures (*n* = 3). **(C)** IL-1β secretion level in human SGC cultures after transfection of WT or mutant GPR37L1 and the effects of paclitaxel (PTX, 1 μg/ml, 24 hours). n = 8 cultures. (**D**) Ti^+^ influx activity is reduced in E296K mutation in human SGCs in the presence of 30 nM MaR1. *n* = 12 cultures. Data are expressed as mean ± SEM, One sample t-test (A), Two-Way ANOVA with Tukey’s post-hoc test (B, C), and One-Way ANOVA with Tukey’s post-hoc test (D) \**P*<0.05, \*\**P*<0.01, \*\*\**P*<0.001.

**Supplemental Figure 10.**
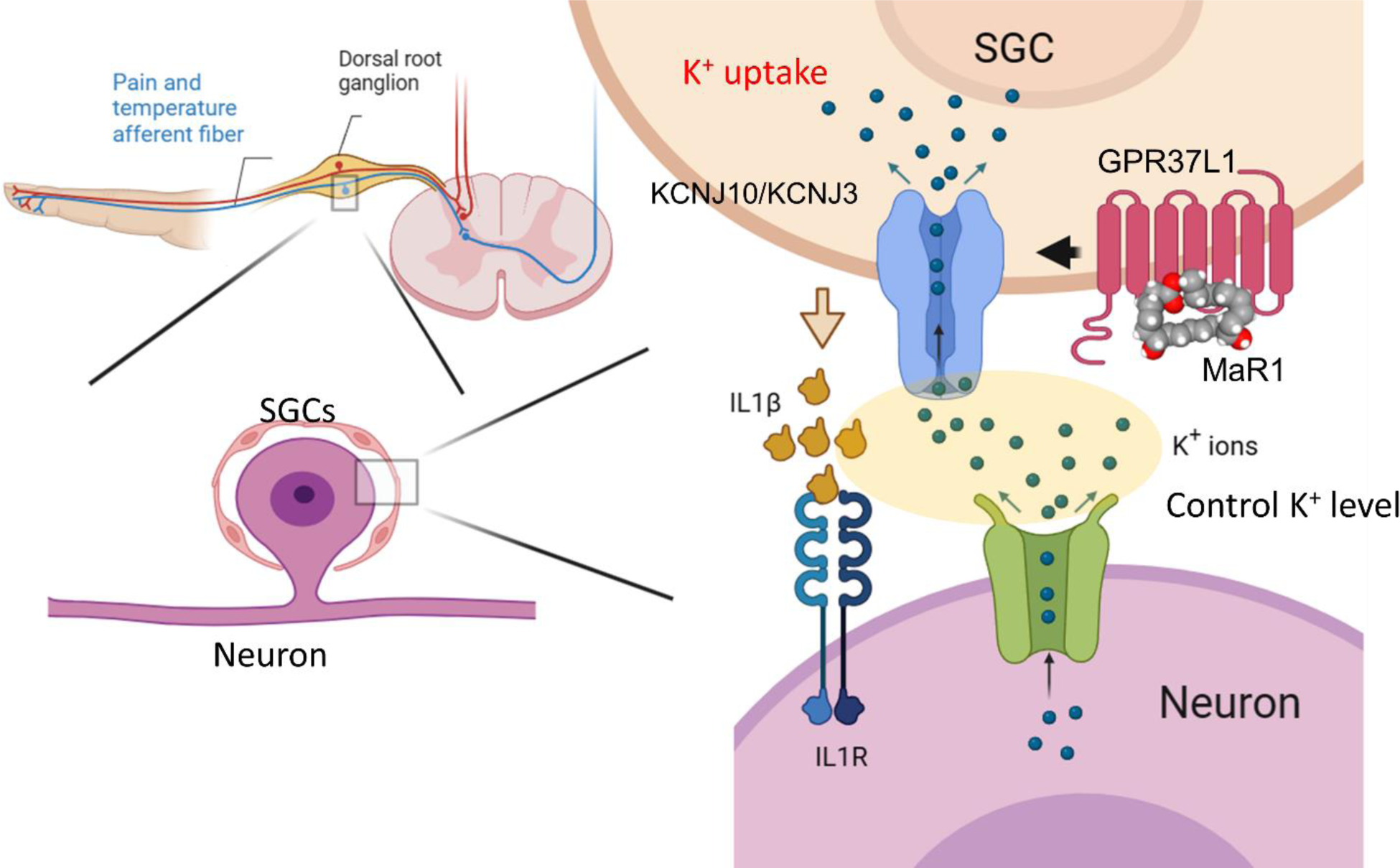
Schematic illustration of GPR37L1-mediated regulation of K^+^ channels in SGCs of DRG. Top left, pain transduction and transmission in peripheral and central axons of DRG neurons. Bottom left, SGCs surrounding a DRG neuron. Right, control of potassium channels signaling by GPR37L1 in SGCs. Neuronal excitation causes K^+^ efflux, and extracellular K^+^ can be up-taken by KCNJ10 in mouse SGCs and KCNJ3/KCNJ10 in human SGCs. Loss of K^+^ uptake activity after chemotherapy and diabetes, as a result of downregulations of GPR37L1/KCNJ3/KCNJ10, will lead to neuropathic pain (CIPN and DPN). Additionally, reduced intracellular K+ levels might initiate ROS-mediated inflammatory responses and contribute to IL-1β production. IL-1β production by both SGCs and neurons can subsequently elevate neuronal excitability (32). Activation of GPR37L1 by MaR1 has the potential to enhance GPR37L1 activity, augment K+ uptake in SGCs, and decrease IL-1β production. This cascade of events may ultimately alleviate pain.

